# RNA polymerase II processing facilitates DNA repair and prevents DNA damage-induced neuronal and developmental failure

**DOI:** 10.1101/2025.03.21.644538

**Authors:** Melanie van der Woude, Karen L. Thijssen, Mariangela Sabatella, Jurgen A. Marteijn, Wim Vermeulen, Hannes Lans

## Abstract

Hereditary transcription-coupled nucleotide excision repair (TC-NER) defects cause severe developmental and neurodegenerative features, as observed in Cockayne syndrome (CS), or mild cutaneous UV sensitivity, as observed in UV-sensitive syndrome. The mechanisms underlying the strikingly different clinical features of these syndromes are not fully understood. Using *C. elegans*, we demonstrate that TC-NER deficiency leads to DNA damage-induced motoneuronal and developmental failure, primarily caused by the lack of lesion removal due to persistent lesion-stalling of RNA polymerase II. If, in the absence of TC-NER, lesion-stalled RNA polymerase II is processed and removed, global genome NER acts as backup pathway to repair transcription-blocking lesions and prevents DNA damage-induced developmental failure. Our results furthermore show that processing of lesion-stalled RNA Polymerase II facilitates TC-NER and involves the activity of multiple E3 ubiquitin ligases. These findings reveal that persistently stalled RNA polymerase II, rather than TC-NER deficiency, is the major driver of severe disease features associated with TC-NER defects.

## Introduction

Accurate transcription of DNA into mRNA by RNA polymerase II (Pol II) is a fundamental requirement for the proper functioning of cells. However, transcription is constantly challenged by DNA damage, which hinders the forward progression of Pol II along the DNA. DNA damage is unavoidable, as cells are continually exposed to environmental and metabolism-derived genotoxic agents, such as UV irradiation, endogenous reactive oxygen species, and aldehydes^1–3^. Different types of DNA damage interfere with transcription by physically blocking Pol II elongation, which may become even more harmful as local Pol II blockage can additionally lead to R-loop and DNA break formation, transcription-replication conflicts, global transcription shutdown, and loss of transcription fidelity^4,5^. This DNA damage-induced transcription stress significantly hampers cellular function, causing cell death and senescence, and is therefore considered to be a major driver of aging^6,7^. To secure transcriptional integrity, and thereby ensure a proper cellular response to stressors, cells eliminate DNA damage in genes via transcription-coupled DNA repair processes, most prominently via transcription-coupled nucleotide excision repair (TC-NER)^4,8–10^.

Nucleotide excision repair (NER) is a major DNA repair pathway which repairs a wide variety of helix-distorting DNA lesions, including those induced by UV light^11,12^. DNA damage can be detected by the global genome NER (GG-NER) subpathway anywhere in the genome or by the TC-NER subpathway upon blockage of elongating Pol II. Detection of lesions by GG-NER relies on the XPC protein, which forms a heterotrimer with CETN2 and RAD23B^13,14^, and constantly probes the DNA searching for helix-distorting DNA lesions^15,16^. Upon stable binding of XPC to a DNA lesion, the core NER pathway is activated by the recruitment of the TFIIH complex ^17,18^. In comparison, detection of lesions by TC-NER relies on activity of the CSB protein, which stably associates with elongating Pol II when it is blocked by a lesion^19,20^. CSB is an ATP-dependent translocase that tries to push Pol II forward over the lesion^21^. If unsuccessful, its stable association with Pol II leads to the recruitment of the E3 ubiquitin ligase complex CRL4^CSA^, consisting of CSA, DDB1, DDA1, RBX1 and CUL4A^22–24^, and, with the help of the Pol II-associated ELOF1 protein^25,26^, to the recruitment of additional TC-NER factors UVSSA and its binding partner USP7^27–31^. During this process, the CRL4^CSA^ complex ubiquitylates both CSB^32^ and Pol II^33–35^, the latter which also involves other E3 ubiquitin ligase complexes^36^, which may lead to their degradation. To promote TC-NER complex stability, the ubiquitin specific protease USP7 deubiquitylates CSB and thereby averts its degradation^27,37^. In addition, ubiquitylation of the RPB1 subunit of Pol II at residue K1268 facilitates the binding of UVSSA, which mediates the recruitment and association of the TFIIH complex to lesion-stalled Pol II^30,33^. GG-NER and TC-NER both merge into a common core NER mechanism, during which TFIIH, with the help of XPA, verifies the lesion^38^. Subsequently, the endonucleases ERCC1-XPF and XPG excise the damaged DNA segment spanning 22-30 base pairs^11,12^ and the resulting gap is filled in by DNA synthesis mediated by replication factors, DNA polymerases and ligases. Once DNA damage is repaired, transcription can restart.

Hereditary TC-NER deficiency manifests clinically as two strikingly divergent syndromes, i.e. Cockayne syndrome (CS) and UV-sensitive syndrome (UV^S^S). CS patients exhibit a range of developmental, progressive neurodegenerative and progeroid features, including early growth cessation, microcephaly, mental retardation with dysmyelination, retinal degeneration, sensorineural deafness, cachexia, photo-hypersensitivity, and a significantly reduced life expectancy^39,40^. In contrast, UV^S^S patients manifest merely mild cutaneous phenotypes, such as photo-hypersensitivity and freckling, without the severe developmental and neurological features seen in CS^31,41,42^. The genetic distinction between both syndromes is mostly found in the affected genes, as *CSA* and *CSB* mutations predominantly lead to CS^43^, while *UVSSA* mutations exclusively lead to UV^S^S^27,28,31^. Few cases are known in which also mutations in *CSA* or *CSB* cause UV^S^S^28,29,31,44,45^. The prominent phenotypic differences have therefore been attributed to additional functions of the CSB and CSA proteins outside TC-NER. For instance, CSB has been implicated in regulating transcription during neuronal differentiation^46^ and in regulating metabolic and DNA repair responses to oxidative stress^47–50^. However, the roles of CSA and UVSSA in several of these processes remain elusive.

The progressive nature of many CS symptoms and the fact that hereditary mutations in downstream NER factors TFIIH, ERCC1-XPF and XPG can cause CS-like features as well^51^, strongly suggest that accumulating DNA damage, not repaired in the absence of TC-NER, is the primary factor underlying these severe CS symptoms. We and others hypothesized that not the DNA damage itself, but the persistent DNA damage-induced transcription stress, characterized by prolonged Pol II stalling at accumulating unrepaired DNA lesions, is the major culprit^4,30,31,33,52–54^. Indeed, CSB and CSA deficient cells are distinguished from UVSSA deficient cells by prolonged chromatin retention of Pol II after DNA damage induction^55,56^. Therefore, it is important to understand the biological impact of prolonged DNA damage-induced Pol II stalling, as this will help to understand the difference in pathogenesis between CS and UV^s^S. During TC-NER, Pol II is likely displaced or even removed so that the downstream repair machinery can gain access to the DNA damage^54,57,58^. In addition, Pol II is thought to be removed from damaged DNA as a last resort pathway alternative to TC-NER, to allow other DNA repair pathways to detect the DNA damage^36^. Pol II processing involves the poly-ubiquitylation and degradation of its major catalytic RPB1 subunit, but many of the mechanistic details are still not fully understood as both CSB and CRL4^CSA^-dependent and -independent mechanisms and multiple additional E3 ubiquitin ligases have been implicated in this processing^33,34,52,59–63^. Impaired Pol II removal will likely shield DNA lesions and prevent them from being repaired via other DNA repair pathways. However, it has not been experimentally shown that other DNA repair pathways can indeed repair transcription-blocking DNA damage if lesion-stalled Pol II is removed. Also, it is unclear if indeed the accumulation of persistent DNA repair intermediates, like lesion-stalled Pol II, is more cytotoxic than the DNA lesions themselves.

The nematode *C. elegans* has become a prominent *in vivo* model organism to investigate diverse aspects of the DNA damage response, including NER, whose mechanism and biological function is highly conserved^64,65^. Previously, we showed that GG-NER is mainly active in the *C. elegans* germ line to preserve the integrity of the entire genome for the next generation. GG-NER deficiency therefore results in meiotic maturation and germ line proliferation defects and embryonic lethality in response to UV irradiation^66,67^. TC-NER, on the other hand, is noticeably active in differentiated, post-mitotic somatic cells in which only the functional integrity of active genes needs to be maintained. We and others showed that, as a consequence, loss of TC-NER function results in strong developmental arrest upon UV irradiation^25,67–70^, making *C. elegans* excellently suited to study the biological impact of DNA damage-induced transcription stress. Here, we investigated the impact of Pol II stalling and the role of various TC-NER proteins during development of *C. elegans* upon DNA damage-induced transcription stress. Our results indicate that this transcription stress causes developmental arrest in TC-NER deficient animals due to persistent Pol II stalling, which is prevented if Pol II is removed and DNA repair by the alternative GG-NER pathway is permitted.

## Results

### DNA damage-induced developmental and neuromotor failure in TC-NER mutants does not always correlate with TC-NER deficiency

To study the biological impact of DNA damage-induced transcription stress in living animals, we determined UV-induced developmental arrest of *C. elegans* larvae with loss-of-function mutations in various TC-NER genes (Table 1), using an L1 larvae UV survival assay^71^. This assay assesses the effect of UV-induced DNA damage on the animal’s ability to develop from first stage (L1) larvae into adults. Importantly, the vast majority of cells in L1 larvae are terminally differentiated^72^, because of which growth and development depend on accurate and intact transcription. We obtained some *C. elegans* loss-of-function mutants from the *Caenorhabditis* Genetics Center and the National Bioresource Project for the nematode, and some were already described by us before (Supplementary Table 1). In line with previous studies^67,69,71,73^, we observed that mutation of the core TC-NER factors *csa-1* (allele *tm5232*) and *csb-1* (allele *ok2335*) results in UV hypersensitivity (Fig. 1A). In comparison, mutation of the core GG-NER factor *xpc-1* (allele *tm3886*) did not affect UV survival. Also, mutation of other TC-NER factors, such as *elof-1* and *math-33*, the *C. elegans* orthologs of mammalian *ELOF1*^25^ and *USP7*^74^, respectively, resulted in clear hypersensitivity and developmental arrest upon UV irradiation (Fig. 1B). These results confirm that the activity of TC-NER proteins CSB-1, CSA-1, ELOF-1 and MATH-33 is essential in somatic tissues of L1 larvae to overcome the cytotoxic consequences of DNA damage-induced transcription stress.

**Table 1.**
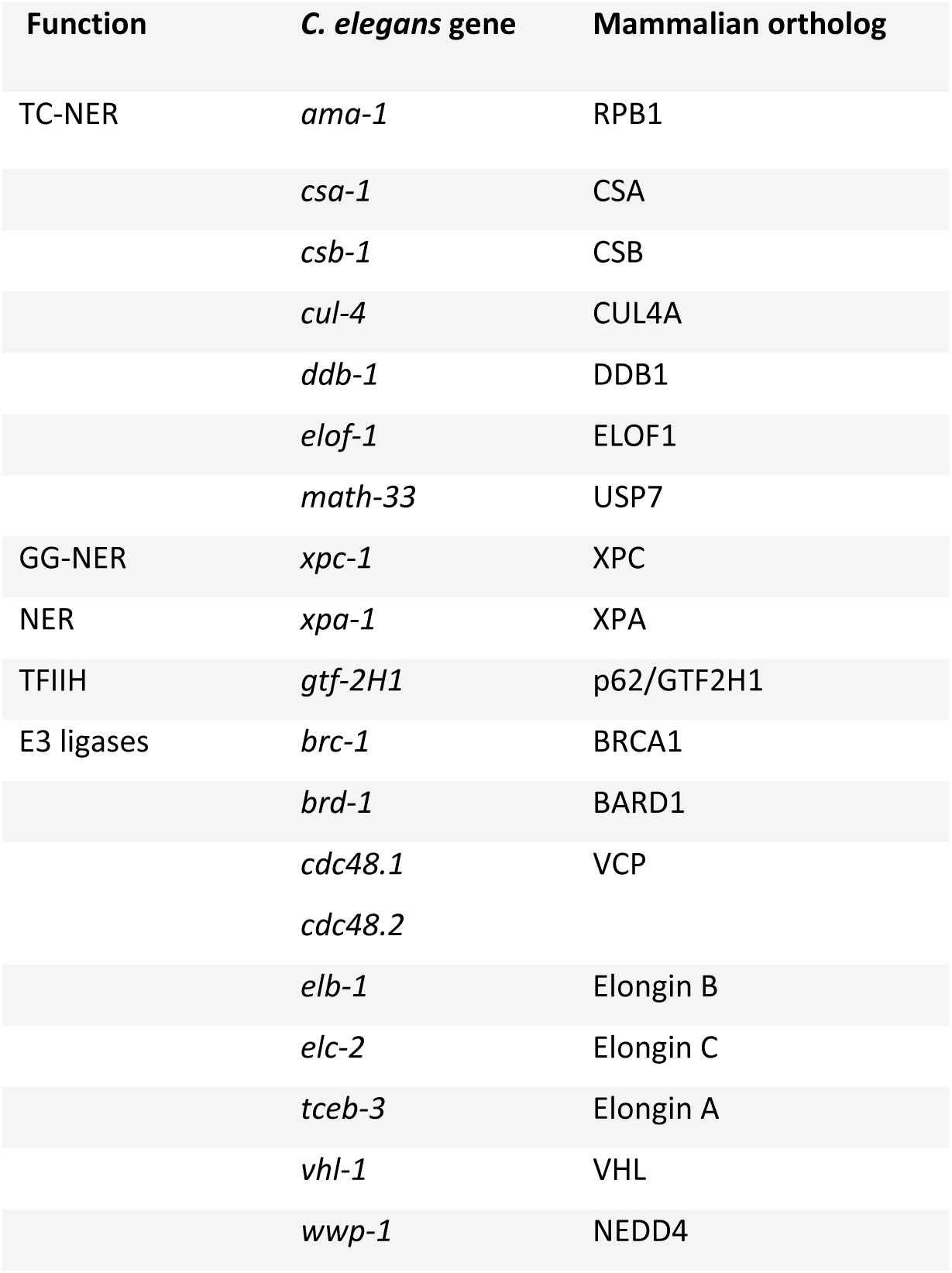
*C. elegans* TC-NER genes studied with their mammalian orthologs.

**Fig. 1.**
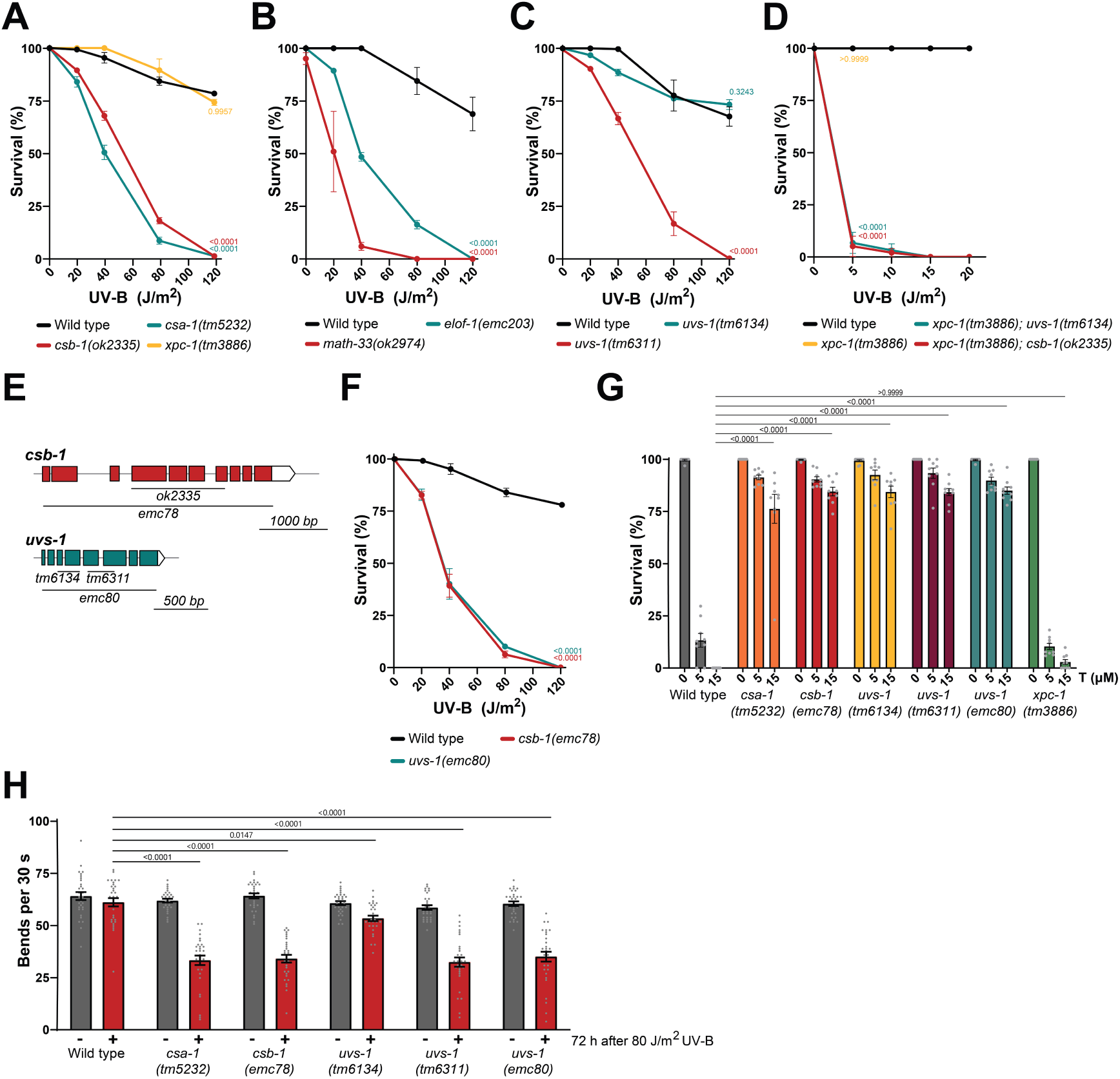
Most but not all TC-NER mutants show developmental and neuromotor failure after DNA damage. (**A-D**) L1 larvae UV survival assays of wild type and (**A**) *csa-1(tm5232)*, *csb-1(ok2335), xpc-1(tm3886);* (**B**) *elof-1(emc203), math-33(ok2974);* (**C**) *uvs-1(tm6134), uvs-1(tm6311);* (**D**) *xpc-1(tm3886); uvs-1(tm6134), xpc-1(tm3886); csb-1(ok2335),* and *xpc-1(tm3886)* animals. (**E**) Schematic depiction of the *csb-1* and *uvs-1* loci with deletion alleles indicated. (**F**) L1 larvae UV survival assay of *csb-1(emc78)* and *uvs-1(emc80)* mutants. (**G**) Trabectedin survival depicting percentage L1 animals that survive beyond the L2 stage of wild type and TC-NER deficient *csa-1(tm5232)*, *csb-1(emc78), uvs-1(tm6134), uvs-1(tm6311),* and *uvs-1(emc80),* and GG-NER deficient *xpc-1(tm3886)* animals. (**H**) Body bends per 30 s of wild type, *csa-1(tm5232), csb-1(emc78), uvs-1(tm6134)*, *uvs-1(tm6311),* and *uvs-1(emc80)* animals, without or 72 h after 40 J/m^2^ UV-B irradiation. Mean with SEM of three independent experiments or (**H**) of two independent experiments. Numbers in the graph represent *p-values* comparing 120 J/m^2^ data points to wild type or as indicated, determined by TWO-WAY ANOVA Šídák’s multiple comparisons test.

Subsequently, we tested UV sensitivity of animals with mutations in the core TC-NER factor *uvs-1*, the nematode ortholog of mammalian *UVSSA*^73^. Strikingly, we observed that the two different mutants that we tested, *uvs-1(tm6134*) and *uvs-1(tm6311)*, showed a distinct developmental arrest upon UV irradiation (Fig. 1C). Whereas *uvs-1(tm6311)* mutants showed a UV hypersensitivity comparable to that of other TC-NER mutants, *uvs-1(tm6134)* mutants were not UV hypersensitive and showed a UV response comparable to that of wild type animals (Fig. 1A, B). This could indicate that *uvs-1(tm6134)* is not a loss-of-function mutant and therefore not TC-NER deficient, while *uvs-1(tm6311)* is. To test this, we crossed the non-UV-hypersensitive *uvs-1(tm6134)* mutant with GG-NER deficient *xpc-1(tm3886)* animals, as we have previously demonstrated that GG-NER acts as backup pathway to TC-NER in repairing DNA damage in somatic tissue^67–69,75^. As a result, GG-NER deficiency does not lead to UV hypersensitivity in TC-NER proficient L1 larvae, but only in TC-NER deficient L1 larvae. Indeed, as shown before^67^, *xpc-1(tm3886); csb-1(ok2335)* double mutant animals showed extreme UV hypersensitivity, even at very low UV doses (Fig. 1D). Strikingly, we found that *xpc-1(tm3886); uvs-1(tm6134)* double mutant animals were similarly extremely UV-hypersensitive (Fig. 1D). This strongly suggests that *uvs-1(tm6134)* animals are TC-NER deficient, despite the lack of clear UV-induced developmental arrest in the single mutant.

The *csb-1(ok2335)* allele is a 1620 bp deletion in exons 4 to 7 that is probably a functional null allele, as it is predicted to encode a truncated CSB-1 protein with most of its catalytic SNF2-like ATPase domain deleted (Fig. 1E)^67^. The *uvs-1(tm6134)* and *uvs-1(tm6311)* alleles are also deletions in exons (Fig. 1E)^73^, that are predicted to encode truncated proteins, as described in more detail below. We therefore tested if the functionality and UV sensitivity of these alleles is comparable to the complete absence of CSB-1 and UVS-1 proteins. To this end, we generated strains with full knock-out alleles of *csb-1* and *uvs-1*, by completely deleting the genes, respectively called *emc78* and *emc80* (Fig. 1E, Supplementary Fig. 1). We observed similar UV-induced developmental arrest for these full knock-out *csb-1(emc78)* and *uvs-1(emc80)* alleles (Fig. 1F) as for the partial gene deletion *csb-1(ok2335)* and *uvs-1(tm6311)* alleles (Fig. 1A,C), respectively.

To determine whether the different *csb-1, csa-1* and *uvs-1* mutants are indeed all TC-NER deficient, we tested their resistance to trabectedin by monitoring larval development after exposure to this drug. Trabectedin, also known as ET743, is an alkylating agent that reacts with guanine residues in the minor groove of the DNA, thereby forming bulky DNA adducts that impede transcription and are recognized by TC-NER ^76^. Trabectedin exposure, however, results in toxicity in TC-NER proficient cells, due to the TC-NER-mediated processing of trabectedin adducts, which leads to the formation of lethal DNA breaks ^77,78^. Therefore, TC-NER deficient cells are resistant to trabectedin exposure. Indeed, we observed that *csa-1(tm5232)* and *csb-1(emc78)* animals were resistant to trabectedin, while wild type and *xpc-1(tm3886)* animals were hypersensitive (Fig. 1G). Furthermore, *uvs-1(tm6311), uvs-1(tm6134)* and *uvs-1(emc80)* mutants were also resistant to trabectedin, which confirms that all three *uvs-1* alleles are true loss-of-function alleles and lead to TC-NER deficiency.

Neuromotor difficulties and neurodegeneration are profound clinical features in TC-NER deficient CS patients. In *C. elegans,* developmental arrest, as measured in the L1 larvae UV survival assay, specifically results from DNA damage interfering with transcription in neurons. We previously showed that intact NER in neurons is sufficient to prevent DNA damage-induced developmental arrest in animals that are otherwise completely NER deficient^68^. To demonstrate this neuronal impact in another manner, we assessed how UV-induced DNA damage affects neuromotor function in TC-NER deficient animals, by determining the total number of lateral body movements per time unit^79^ before and after UV irradiation (Fig. 1H). Similar to what was observed in the L1 larvae UV survival assay, *csb-1(emc78)*, *csa-1(tm5232)*, *uvs-1(emc80)*, and *uvs-1(tm6311)* were UV hypersensitive, as they displayed significantly impaired motility after UV-induced DNA damage. In contrast, *uvs-1(tm6134)* mutant animals again did not display any UV hypersensitivity, as their motility after UV-induced DNA damage was comparable to that of wild type animals. Taken together, these findings show that UV-induced DNA damage leads to both developmental arrest and neuromotor difficulties in most TC-NER deficient animals. However, the absence of UV hypersensitivity in TC-NER deficient *uvs-1(tm6134)* animals suggests that these phenotypes are not primarily due to the TC-NER deficiency per se, but likely due to an alternative cause.

### Deficient USP7 and TFIIH recruitment cause UV hypersensitivity in *uvs-1* mutants

To understand the difference in UV sensitivity between the *uvs-1(tm6134)* and *uvs-1(tm6311)* alleles, we tested how these alleles affect the expression and protein levels of UVS-1 and CSB-1. The mammalian UVS-1 ortholog UVSSA contains several domains that are important for its function within the TC-NER complex, including a VHS domain that mediates interaction with CSA^29,30,35,80^, a TRAF-binding motif that mediates interaction with USP7^81^, a p62/GTF2H1-binding region^82^, and a DUF2043 domain that plays an important role in its recruitment to DNA damage^80^. We mapped these UVSSA protein domains to the *C. elegans* UVS-1 protein sequence, showing that UVS-1 also contains these essential TC-NER interaction domains (Fig. 2A; Supplementary Fig. 2). Next, we assessed which domains are affected by the *tm6134* and *tm6311* alleles. As both *tm6134* and *tm6311* deletions span multiple exons and introns, we first confirmed by RT-PCR that both mutant alleles are expressed (Fig. 2B). Subsequently, we sequenced cDNA of both alleles, to determine the ATG start site and how the genomic deletions affect splicing of the mRNA (Supplementary Fig. 3). Based on this sequence, protein prediction according to FGENESH^83^ suggested that the non-UV-hypersensitive *uvs-1(tm6134)* allele encodes a protein in which the VHS domain is partially deleted (Fig. 2A; Supplementary Fig. 2). In comparison, the protein encoded by the UV-hypersensitive *uvs-1(tm6311)* allele still retains the VHS domain, but lacks the TRAF-binding motif and most of the GTF-2H1 binding region, and therefore likely will not be able to bind to the *C. elegans* USP7 ortholog MATH-33 and the p62/GTF2H1 ortholog GTF-2H1, which is a subunit of TFIIH.

**Fig. 2.**
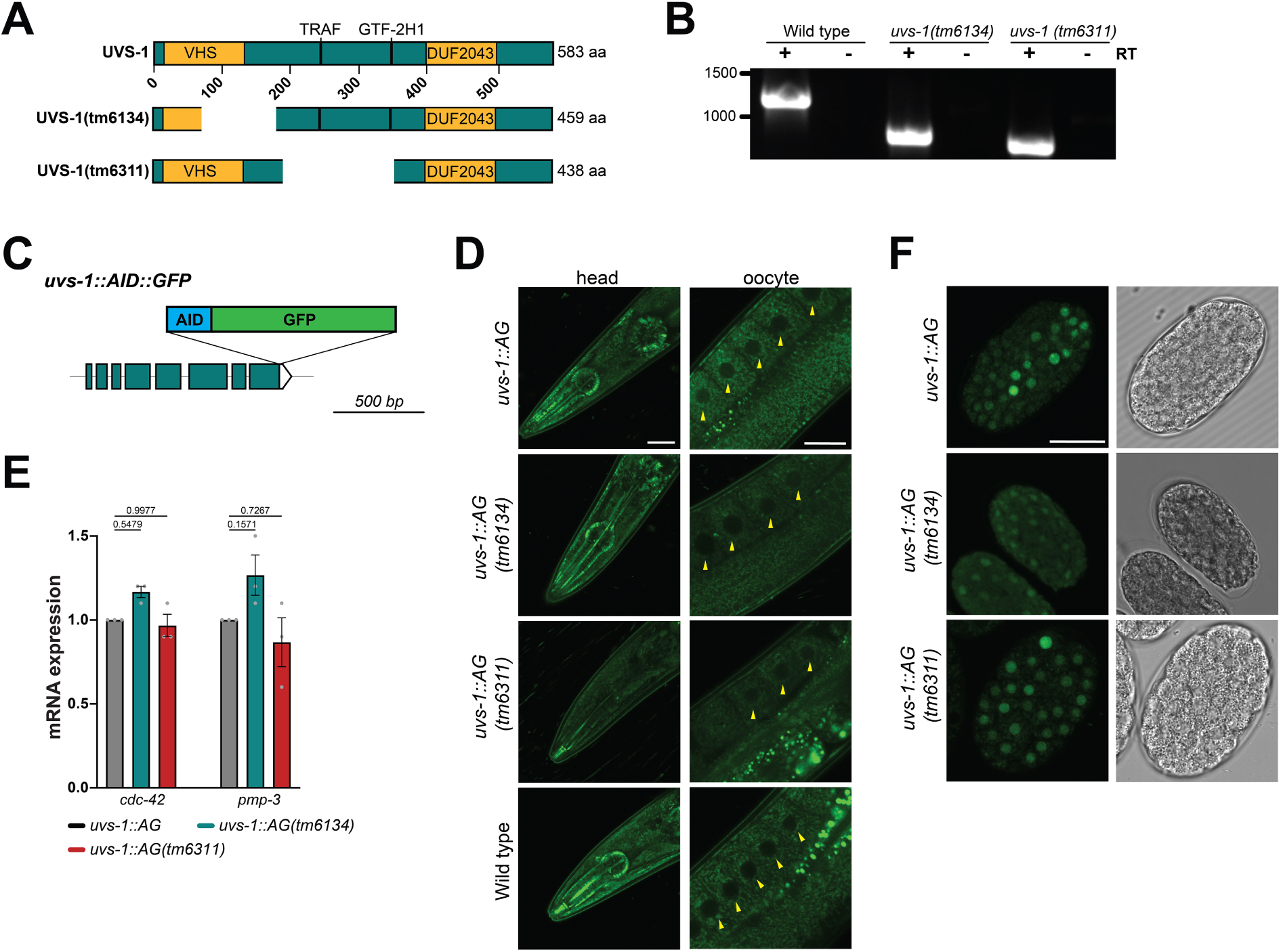
UVS-1 mutants and expression. (**A**) FGENESH prediction of the UVS-1 protein encoded by wild type *uvs-1*, *uvs-1(tm6134)*, and *uvs-1(tm6311)* alleles. (**B**) RT-PCR on exons 1-6 of cDNA of wild type, *uvs-1(tm6134)*, and *uvs-1(tm6311)* alleles. (**C**) Schematic depiction of AID::GFP knock-in at the *uvs-1* locus. (**D**) Representative fluorescence confocal microscopy images of living animals showing no observable UVS-1::AID::GFP expression in head and oocyte nuclei of *uvs-1::AG* knock-in wild type, *uvs-1(tm6134)*, and *uvs-1(tm6134)* animals. Wild type animals, not expressing GFP (lower panel), are included as reference for autofluorescence signals. Arrowheads indicate nuclei. Scale bar: 25 µm. (**E**) Relative mRNA levels as determined by qPCR of *uvs-1::AG, uvs-1::AG(tm6134*) and *uvs-1::AG(tm6311)* alleles. Numbers in the graph represent *p-values* compared to wild type. Results are normalized to control household genes *pmp-3* or *cdc-42*, as indicated, and plotted as mean with SEM of three independent experiments. (**F**) Representative confocal images of animals overexpressing wild type or mutant UVS-1::AG in embryonic nuclei. Right panels show brightfield images. Scale bar: 20 µm

To test if the mutant alleles express truncated UVS-1 proteins and whether there are differences in the expression and tissue distribution of wild type and mutant UVS-1, we knocked in GFP fused to an auxin-inducible degradation tag (AID::GFP; abbreviated to ‘AG’ for simplicity) at the C-terminus of wild type *uvs-1* and the *uvs-1(tm6134)* and *uvs-1(tm6311)* alleles (Fig. 2C). Functionality of the tagged UVS-1 proteins was confirmed by L1 larvae UV survival experiments, showing that the UV survival of the wild type and *tm6134* mutant knock-in animals expressing AG-tagged UVS-1 was similar as that of wild type animals (Supplementary Fig. 4). This clearly indicates that UVS-1::AG must be normally expressed and biologically active. Nevertheless, using confocal microscopy, we could not detect any expression of UVS-1::AG in nuclei of different tissues, such as in the head and in oocytes (Fig. 2D). Note that the fluorescence visible in the images of Fig. 2D is not derived from the GFP tag but from non-specific background autofluorescence, as this was also observed in animals without a GFP tag. In contrast, previously generated similar knock-in strains for other NER factors, i.e. XPF-1 and TFIIH (GTF-2H1 subunit), showed clear nuclear expression and ubiquitous distribution of these proteins^68,84^ (Supplementary Fig. 5). Also, no UVS-1::AG expression was detected in the *uvs-1::AG(tm6134)* and *uvs-1::AG(tm6311)* knock-in strains. We confirmed that all three knock-in alleles are expressed, using RT-qPCR on GFP (Fig. 2E). This showed that *uvs-1::AG(tm6134)* and *uvs-1::AG(tm6311)* mRNA levels are similar as those of wild type *uvs-1::AG*, ruling out that the differences observed between UV-induced developmental and neuronal impairment are caused by differences in UVS-1 expression levels. As the expression levels of *uvs-1::AG* are apparently below the detection limit of our employed confocal microscope, we generated animals in which the same AG-tagged wild type and mutant UVS-1 genes driven by the *uvs-1* promoter are overexpressed. In these animals, nuclear expression of both wild type and mutant UVS-1::AG was clearly visible (Fig. 2F).

To test whether other TC-NER proteins are expressed similarly at low levels, we determined the expression of CSB-1, by generating *csb-1::AID::mScarlet3* knock-in animals (Fig. 3A). We chose mScarlet3, as this is the brightest red fluorescent protein currently available^85^ and red fluorescence imaging generates lower autofluorescence levels than green fluorescence imaging in *C. elegans*^86^. Normal expression and functionality of the tagged CSB-1 protein was confirmed by the L1 larvae UV survival assay (Supplementary Fig. 4), but again this expression was below the detection limit of our confocal microscope in various somatic tissues of living animals, including in the head (Fig. 3B). Strikingly, clear expression of CSB-1::AID::mScarlet3 was observed, but only in nuclei of meiotic cells and oocytes of the germline and in embryos. Even though CSB-1 expression could not be detected in differentiated somatic tissues by microscopy, previous microarray gene expression analyses have shown that CSB-1 is expressed in these tissues^87,88^. Moreover, we and others showed that mutation of *csb-1* impairs DNA repair, survival and functionality of neurons and muscle cells in *C. elegans*, clearly indicating that CSB-1 is expressed and active in these tissues^68,75,87,89–91^. To confirm this, we generated animals overexpressing FLAG::GFP-tagged CSB-1 from its own promoter, which indeed showed clear expression of CSB-1 in somatic cells (Fig. 3C).

**Fig. 3.**
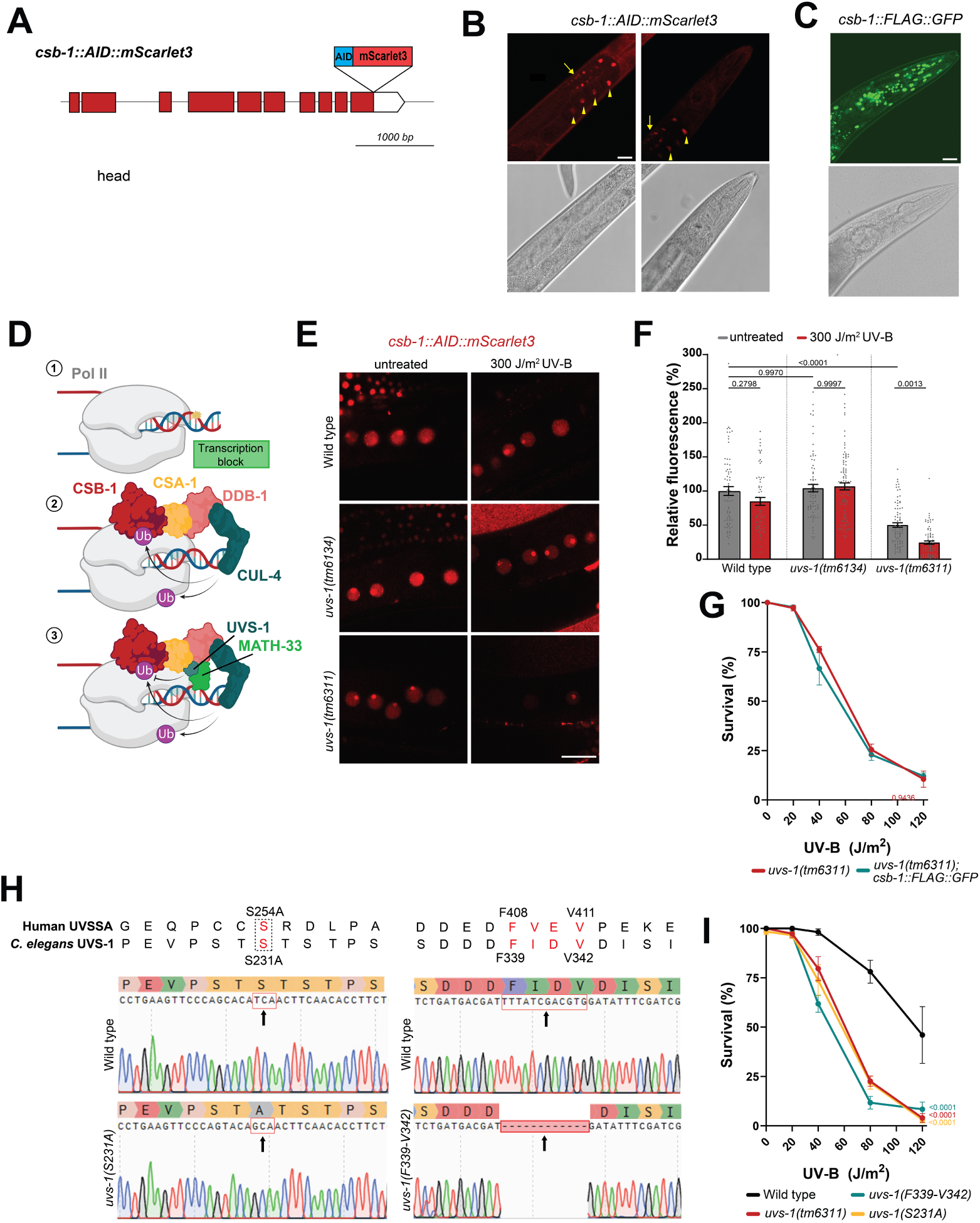
Deficient USP7 and TFIIH recruitment cause UV hypersensitivity in *uvs-1* mutants. (**A**) Schematic depiction of AID::mScarlet3 knock-in at the *csb-1* locus. (**B**) Representative confocal images of fixed animals showing CSB-1::AID::mScarlet3 expression in pachytene (arrow) and oocyte (arrowheads) nuclei in the germline (left panel) and no observable expression in head nuclei (right panel). Lower panel shows brightfield images. Scale bar: 20 µm (**C**) Representative confocal image of animals overexpressing CSB-1::FLAG::GFP in somatic nuclei in the head. Lower panel shows a brightfield image. Scale bar: 20 µm (**D**) Schematic depiction of the first step of the TC-NER reaction. TC-NER is initiated when elongating Pol II, including catalytic subunit AMA-1, is stalled by DNA damage in the transcribed strand of an active gene (Step 1). This leads to the stable binding of TC-NER factors CSB-1 and CRL4^CSA-1^, including CSA-1, DDB-1, CUL-4A and RBX-1 (not shown), which ubiquitylates CSB-1 and lesion-stalled Pol II (Step 2). Subsequently, UVS-1 and its binding partner MATH-33 are recruited, which leads to deubiquitylation of CSB-1 by MATH-33 and thereby stabilization of the TC-NER complex (Step 3). (**E-F**) Representative fluorescence images (**E**) and quantification (**F**) of CSB-1::AID::mScarlet3 expression in nuclei of oocytes in wild type, *uvs-1(tm6134)*, and *uvs-1(tm6311)* mutants. Fluorescence values were normalized to CSB-1::AID::mScarlet3 in wild type animals. Mean with SEM of two independent experiments. (**G**) L1 larvae UV survival assays of *uvs-1(tm6311)* animals without and with overexpression of CSB-1::FLAG::GFP. Mean with SEM of three independent experiments. (**H**) Genomic and predicted protein sequences of the wild type and S231A and F339-V342 mutant *uvs-1* alleles, compared to the mammalian ortholog UVSSA sequences. (**I**) L1 larvae UV survival assays of wild type, *uvs-1(tm6311)*, *uvs-1(F339-V342)* and *uvs-1(S231A)* animals. Mean with SEM of three independent experiments. Numbers in the graph represent *p-values* comparing the 120 J/m^2^ data points to wild type, determined by TWO-WAY ANOVA Šídák’s multiple comparisons test.

A major difference between the proteins encoded by both mutant *uvs-1* alleles is that UVS-1(tm6311) is predicted to lose its interaction with MATH-33/USP7 due to deletion of the TRAF-binding motif, while UVS-1(tm6134) still retains this interaction. To test the importance of this interaction, we crossed *uvs-1(tm6134)* mutants with *math-33* mutants and observed that *uvs-1(tm6134)* animals do become UV hypersensitive upon MATH-33/USP7 loss (Supplementary Fig 4B). In humans, the deubiquitylase USP7 is recruited to TC-NER complexes through this interaction with UVSSA to counteract the degradation of CSB after its ubiquitylation by CRL4^CSA^, thereby stabilizing the TC-NER complex^27,28,37,92^ (Fig. 3D). We therefore hypothesized that *C. elegans* CSB-1 protein levels could be affected by the *tm6311* deletion in UVS-1. We crossed both *uvs-1* mutants with *csb-1::AID::mScarlet3* knock-in animals to study CSB-1 levels. Quantification of mScarlet3 fluorescence levels in oocytes of living animals by confocal microscopy showed no difference in CSB-1 protein levels between wild type and *uvs-1(tm6134)* animals, in untreated conditions and after UV irradiation (Fig. 3E, F). However, CSB-1 levels were strongly reduced in *uvs-1(tm6311)* mutants, already in untreated conditions and even more so after UV irradiation (Fig. 3E, F). These results indicate that CSB-1 is unstable and degraded in *uvs-1(tm6311)* mutants, likely because it is not deubiquitylated by MATH-33, and that possibly *uvs-1(tm6311)* mutants are UV hypersensitive due to low CSB-1 levels.

We therefore tested if increased CSB-1 levels could rescue the UV hypersensitivity of *uvs-1(tm6311)* animals, by crossing these animals with animals overexpressing GFP-tagged CSB-1, driven by its own promoter. L1 larvae UV survival assays did not show any difference upon CSB overexpression (Fig. 3G), indicating that *uvs-1(tm6311)* mutants are not UV hypersensitive solely due to low CSB-1 levels. The mutant UVS-1(tm6311) protein is not only predicted to have lost its interaction with MATH-33/USP7, but also with the TFIIH subunit GTF-2H1. To test which of these two lost interactions is responsible for the UV hypersensitivity, we generated two additional *uvs-1* mutants (Fig. 3H). In one mutant, we mutated serine residue Ser231 in the TRAF-binding motif of UVS-1 into arginine, which in human UVSSA was shown to lead to loss of the interaction with USP7^81^ In the other mutant, we deleted residues F51 to V54 of the GTF-2H1 binding region, which in human UVSSA was shown lead to loss of GTF2H1 binding ^18^. Interestingly, both mutants were hypersensitive to UV irradiation, similarly as *uvs-1(tm6311)* mutants (Fig. 3I). These results indicate that the interactions of UVSSA both with USP7 and with TFIIH are needed to protect against UV-induced developmental arrest.

### Persistently lesion-stalled Pol II in TC-NER mutants correlates with developmental arrest

In humans, binding of CSB to lesion-stalled Pol II leads to recruitment of the CRL4^CSA^ E3 ubiquitin ligase complex, and possibly other E3 ubiquitin ligases, which are thought to poly-ubiquitylate, besides CSB, also the major catalytic Pol II subunit RPB1. This leads to the displacement and/or removal of the lesion-stalled Pol II complex to allow repair to take place (Fig. 3D)^9,10^. Also, the recruitment of TFIIH was hypothesized to lead to Pol II displacement^4,93–98^. Thus, a key feature of UV hypersensitive *csb-1* and *uvs-1* strains may be that Pol II is not efficiently removed from damaged DNA. In order to study the DNA binding of Pol II in living worms, we created *AID::GFP::3xFLAG::ama-1* (referred to as *AGF::ama-1* for simplicity) animals, in which GFP fused to an AID and a triple FLAG tag is knocked in at the N-terminus of the *ama-1* locus, which encodes the ortholog of the human Pol II subunit RPB1 (Fig. 4A). Importantly, *AGF::ama-1* knock-in animals were fully viable and did not show any UV hypersensitivity compared to wild type animals (Supplementary Fig. 4), indicating that the triple tag does not interfere with Pol II function. Expression of AGF::AMA-1 was clearly observed in all nuclei of each cell type throughout development, in line with the need for transcription in every cell (Fig. 4B), contrasting with the low expression of UVS-1 and CSB-1. Interestingly, AGF::AMA-1 fluorescence was more than five times higher in intensity compared to TFIIH subunit GTF-2H1 tagged with AID::GFP (Supplementary Fig. 5)^84^, indicating that AMA-1 is expressed at much higher levels than the NER and transcription factor TFIIH. TC-NER factors, despite their importance to DNA repair, are apparently expressed at even lower levels compared to other NER factors.

**Fig. 4.**
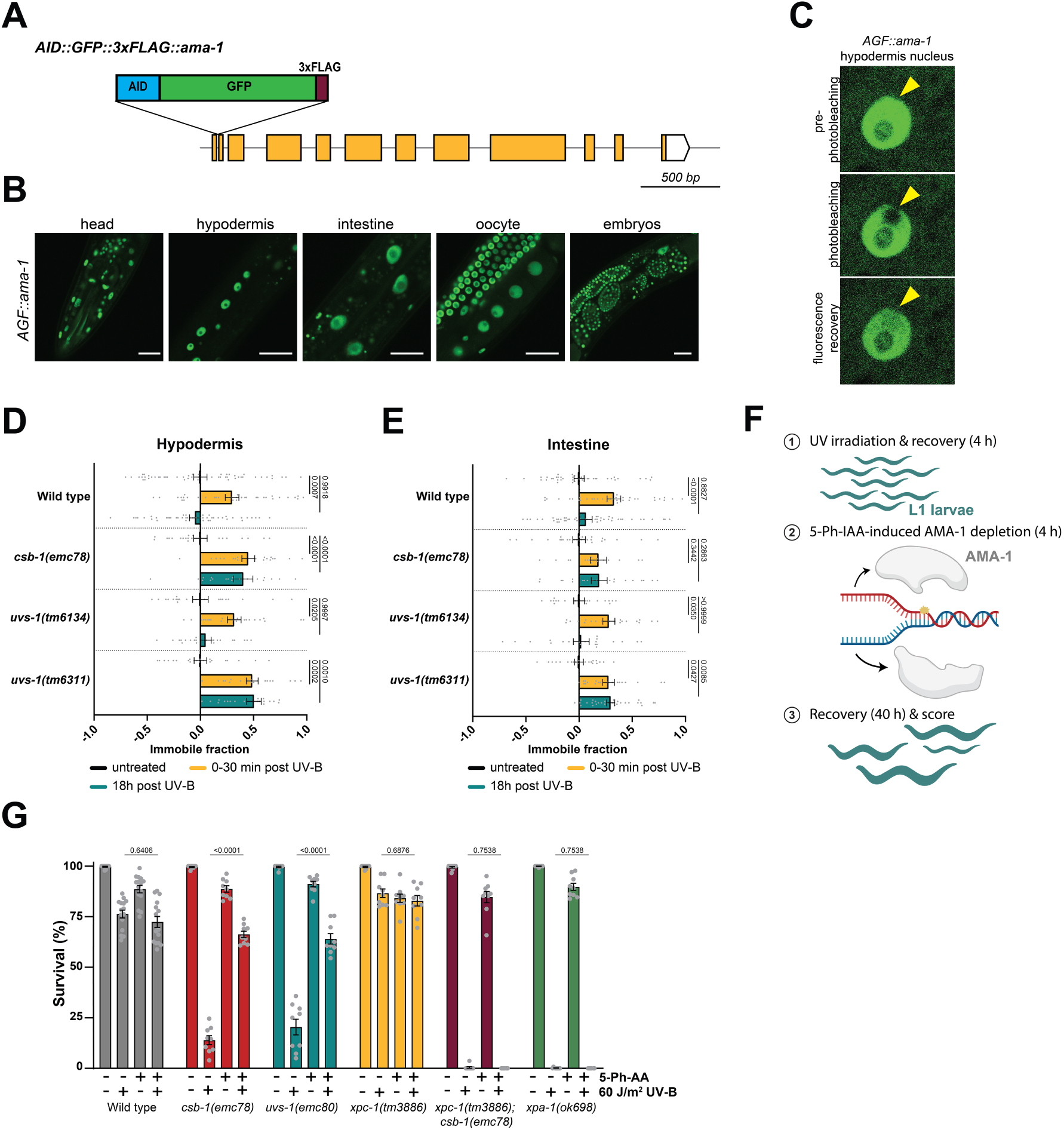
Pol II processing in TC-NER mutants facilitates GG-NER of transcription-blocking lesions. (**A**) Schematic depiction of *AID::3xFLAG::GFP* knock-in at the *ama-1* locus (**B**) Representative fluorescence confocal microscopy images of living animals showing AGF::AMA-1 expression in the head, hypodermis, intestine, oocyte, and embryos. Scale bar: 50 µm. (**C**) Representative FRAP images of a hypodermis nucleus with AGF::AMA-1 expression, before photobleaching (top), after photobleaching (middle) and after recovery of fluorescence. (**D-E**) AGF::AMA-1 immobile fractions, calculated from FRAP curves depicted in Supplementary Fig. 6A-B, performed under untreated conditions, directly (0-30 min) or 18 h after UV-B irradiation, in hypodermal (**D**) or intestinal (**E**) nuclei of wild type, *csb-1(emc78)*, *uvs-1(tm6134)*, and *uvs-1(tm6311)* knock-in animals expressing AGF::AMA-1. Mean with SEM of three independent experiments. (**F**) Experimental set-up of an L1 larvae UV survival assay during which Pol II was transiently depleted. L1 larvae were untreated or irradiated (60 J/m^2^ UV-B) and left to grow for 4 h (Step 1). Next, animals were cultured in the absence or presence of 5-Ph-IAA for 4 h to transiently deplete AGF::AMA-1 (Step 2). After 40 h, animal development wase scored (Step 3). (**G**) L1 larvae UV survival assay after 5-Ph-IAA-induced Pol II depletion in wild type, *csb-1(emc78)*, *uvs-1(emc80), xpa-1(ok698), xpc-1(tm3886)*, and *xpc-1(tm3886); csb-1(emc78)* animals expressing AGF::AMA-1. Mean with SEM of minimal three independent experiments. Numbers in the graph represent *p-values* determined by TWO-WAY ANOVA Šídák’s multiple comparisons test.

To study how *csb-1* and *uvs-1* loss-of-function affect Pol II binding to DNA damage, we crossed *AGF::ama-1* animals with the UV hypersensitive *csb-1(emc-78)* and *uvs-1(tm6311)* and the non-UV hypersensitive *uvs-1(tm6134)* strains. We assessed Pol II mobility and DNA binding using fluorescence recovery after photo-bleaching (FRAP)^99^, in hypodermal and intestinal nuclei of living AGF::AMA-1-expressing animals immediately and 18 h after induction of DNA damage by UV irradiation (Fig. 4C). UV-induced immobilization of Pol II, measured as incomplete fluorescence recovery in FRAP, reflects its stalling at UV-induced DNA lesions and thus its binding to damaged DNA^100^, as previously also shown for human fluorescently-tagged Pol II subunit RPB1^101^. In wild type animals, a significant fraction of AGF::AMA-1 molecules indeed became immobilized within the first 30 min after UV irradiation, in both hypodermal and intestinal cells, as shown by the immobile fraction (Fig. 4D-E) calculated based on the FRAP curves (Supplementary Fig. 6A-B). This immobile fraction was not observed anymore 18 h after UV irradiation, indicating that AMA-1 was no longer bound to damaged DNA due to functional TC-NER and thus proper processing of Pol II. Also, in *csb-1(emc78)* and both *uvs-1* mutants a significant fraction of AGF::AMA-1 became immobilized immediately after DNA damage induction, in both hypodermal and intestinal cells. However, in the UV hypersensitive *csb-1(emc78)* and *uvs-1(tm6311)* mutants a clear UV-induced immobile fraction was still observed 18 h after UV irradiation. In contrast, in the non-UV hypersensitive *uvs-1(tm6134)* strain, this immobile fraction was not observed anymore after 18 h, similar as in wild type animals (Fig. 4D-E; Supplementary Fig. 6A-B). These observations show that Pol II is effectively processed and removed from damaged DNA in wild type and *uvs-1(tm6134)* animals, but is persistently stalled at UV-induced DNA lesions in the UV hypersensitive *csb-1(emc78)* and *uvs-1(tm6311)* strains. Therefore, these results indicate that persistent DNA damage-induced Pol II stalling correlates with developmental arrest in *csb-1(emc78)* and *uvs-1(tm6311)* animals.

### Transient Pol II depletion rescues UV hypersensitivity of *csb-1* and *uvs-1* animals in a GG-NER dependent manner

Persistent Pol II stalling may cause developmental arrest by shielding the lesion from repair via other DNA repair pathways and thus impeding transcription restart. To investigate if indeed the lesion-stalling of Pol II is responsible for the developmental arrest, we tested if removal of Pol II from damaged DNA rescues the UV hypersensitivity of *csb-1(emc78)* and *uvs-1(emc80)* mutants. We therefore made use of the AID degron tag^102^ fused to AMA-1 in *AGF::ama-1* animals to transiently deplete AMA-1, while performing L1 larvae UV survival assays. To this end, we expressed the improved Arabidopsis E3 ubiquitin ligase mutant TIR1(F79G)^103^, under control of the ubiquitously expressed ribosomal protein promoter *rps-28*, in wild type, *csb-1(emc78)* and *uvs-1(emc80)* animals. TIR1(F79G) is activated by culturing animals on the auxin derivative 5-Ph-IAA, leading to depletion of AID-tagged AMA-1 (Supplementary Fig. 6C). To avoid too exhaustive depletion of AGF::AMA-1, which would lead to complete loss of transcription and larval arrest by itself, we cultured AGF::AMA-1 L1 larvae for 4 h only on 5-Ph-IAA during the L1 larvae UV survival assay to transiently deplete Pol II (Fig. 4F). This transient Pol II depletion only minimally affected the development of unirradiated wild type animals and did not lead to any additional developmental arrest after UV irradiation in wild animals (Fig. 4G). Strikingly, however, this transient Pol II depletion significantly rescued the UV hypersensitivity of the TC-NER deficient *csb-1(emc78)* and *uvs-1(emc80)* strains, to a level comparable to that of UV-irradiated wild type animals in which Pol II was depleted (Fig. 4G).

The remarkable reversal in UV hypersensitivity observed in these TC-NER deficient strains suggests that removal of stalled Pol II allows the lesion to become accessible for detection and repair via alternative DNA repair pathways. As the GG-NER pathway is the most likely candidate for the removal of UV-induced lesions, we tested if GG-NER is responsible for the rescued UV hypersensitivity. We crossed the TIR1(F79G)- and AGF::AMA-1-expressing wild type and *csb-1(emc78)* animals with *xpc-1(tm3886)* mutants that are deficient in GG-NER. Also, we crossed the TIR1(F79G)- and AGF::AMA-1-expressing wild type strain with an *xpa-1(ok698)* mutant strain, deficient for the NER factor XPA-1 that is essential for both GG-NER and TC-NER^67,70^. GG-NER deficient *xpc-1(tm3886)* animals showed no hypersensitivity to UV irradiation, irrespective of whether Pol II was depleted, which is as shown before (Fig. 1A)^67^ and consistent with the proficiency of TC-NER in these animals. However, the rescued UV hypersensitivity observed after transient Pol II depletion in TC-NER deficient *csb-1(emc78)* animals was completely abolished when in addition also GG-NER was deficient, i.e. in *xpc-1(tm3886); csb-1(emc78)* double mutants (Fig. 4G). GG-NER and TC-NER deficient *xpa-1(ok698)* mutants similarly showed strong UV hypersensitivity, irrespective of whether Pol II was transiently depleted or not. Together, these findings demonstrate that the shielding and prevention of repair of DNA lesions by persistently stalled Pol II is the primary cause of developmental arrest in UV-irradiated TC-NER deficient *C. elegans* L1 larvae. Importantly, our data show that this developmental arrest can be averted if Pol II is removed from DNA lesions, allowing GG-NER to repair the UV-induced DNA damage.

### Multiple E3 ubiquitin ligases promote Pol II processing to prevent DNA damage-induced developmental arrest

To confirm that the inability to displace and/or remove lesion-stalled Pol II impairs cell function and consequently development, we investigated which other factors, besides CSB-1 and CSA-1, are involved in this Pol II processing. In mammals, RPB1 is poly-ubiquitylated at residue K1268, which involves the activity of CRL4^CSA^ and likely also other E3 ubiquitin ligases^24,33,36^. This ubiquitylation was found to promote both TC-NER and Pol II degradation, as well as transcription recovery in human cells, and to protect against CS-like features, including neurodegeneration, in mice^33,34^. Defective K1268 ubiquitylation was therefore proposed to lead to persistent stalling of Pol II at unrepaired lesions, which may impede access of alternative DNA repair pathways. The RPB1 residue K1268 is well conserved among various species and is in *C. elegans* represented by residue K1260 in *ama-1* (Fig. 5A). We inactivated this potential ubiquitylation site in *C. elegans,* by mutating the lysine at position 1260 to arginine, creating a strain referred to as *ama-1(K1260R)*. Subsequently, we performed L1 larvae UV survival assays, in which we observed similar UV-induced developmental arrest as in the UV hypersensitive TC-NER deficient mutants (Fig. 5B; compare to Fig. 1A-C, F). This finding supports the idea that the UV-induced developmental arrest, as measured in the L1 larvae UV survival assay, is caused by deficient removal of Pol II from DNA damage.

**Fig. 5.**
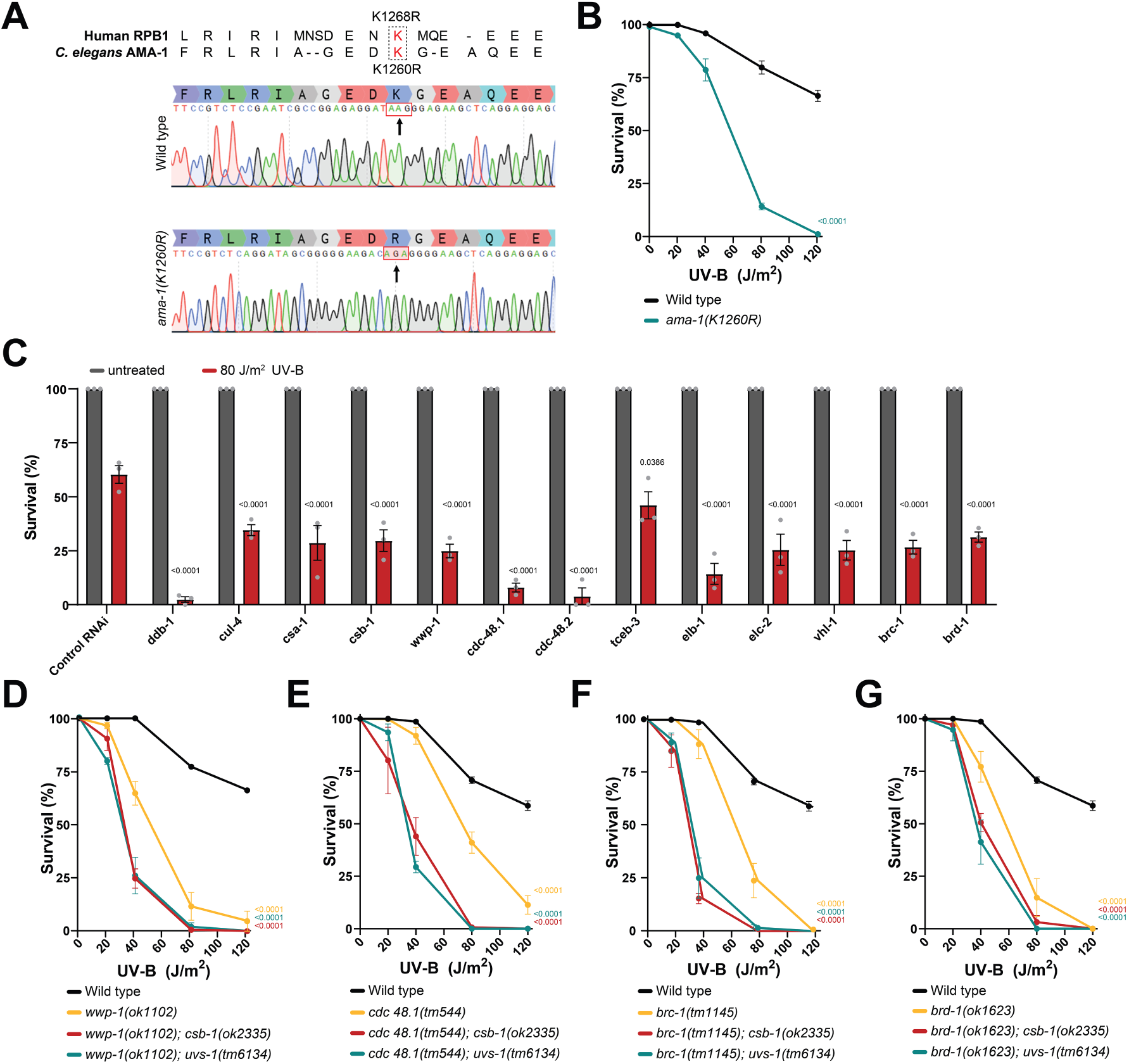
Multiple E3 ubiquitin ligases promote Pol II processing to prevent developmental arrest. (**A**) Genomic and protein sequences of the wild-type and ubiquitylation-defective K1260R mutant *ama-1* allele, compared to its mammalian ortholog RPB1 and corresponding K1286R mutant residue. (**B**) L1 larvae UV survival of wild type and *ama-1(K1260R)* animals. (**C**) L1 larvae UV survival of animals treated with RNAi against TC-NER genes (*ddb-1, cul-4, csa-1,* and *csb-1*), and orthologs of NEDD4 (*wwp-1*), VCP (*cdc-48.1,* and *cdc-48.2*), Elongin-Cullin complexes (*tceb-3, elb-1, elc-1,* and *vhl-1*), and the BRCA1/BARD1 complex (*brc-1* and *brd-1*). (**D-G**) L1 larvae UV survival experiments of wild type and single mutants of (**D**) *wwp-1(ok1102)*, (**E**) *cdc-48.1(tm544)*, (**F**) *brc-1(tm1145)*, and (**G**) *brd-(ok1623)* and of double mutants of each of these genes together with either *csb-1(ok2335)* (red) or *uvs-1(tm6134)* (cyan). All graphs depict mean with SEM of three independent experiments. Numbers in the graph represent *p-values* comparing 80 or 120 J/m^2^ data points to wild type, determined by TWO-WAY ANOVA Šídák’s multiple comparisons test.

Besides CRL4^CSA^, several E3 ubiquitin ligases have been implicated in ubiquitylation and degradation of Pol II stalled at transcription-blocking DNA lesions ^36^. These are in humans NEDD4^59,60^, Elongin-Cullin complexes CRL5^Elongin59,61,104^ and CRL2^VHL62,105^, and BRCA1/BARD1^63,106^. Also, the ubiquitin-dependent segregase VCP was implicated in removing stalled ubiquitylated Pol II from DNA damage^56,107–109^. We tested whether each of these factors is important to prevent UV-induced developmental arrest, by performing L1 larvae UV survival assays after knocking down their *C. elegans* orthologs using RNAi (Fig. 5C; Table 1)^110^. Knockdown of *wwp-1*, the *C. elegans* ortholog of human NEDD4, previously already implicated in ubiquitylating AMA-1^70^, and two genes encoding orthologs of VCP, *cdc-48.1* and *cdc-48.2,* gave rise to UV hypersensitivity, similarly as knockdown of the CRL4^CSA^ complex factors *ddb-1, cul-4, csa-1,* and of *csb-1,* which were included as control (Fig. 5C). Also, knockdown of CRL5^Elongin^ and CRL2^VHL^ complex genes, i.e. *tceb-3*, *elc-2*, *elb-1*, and *vhl-1*, which are the orthologs of human Elongin A, Elongin B, Elongin C, and VHL, respectively, led to UV hypersensitivity. This was also observed after knockdown of the BRCA1 and BARD1 orthologs *brc-1* and *brd-1*. These findings corroborate the idea that Pol II processing is necessary to prevent DNA damage-induced developmental arrest and suggest that this is an intricate process involving the activity of multiple E3 ubiquitin ligases.

To further confirm these findings, we obtained loss-of-function mutants for several of the tested genes, i.e. *wwp-1(ok1102)*, *cdc-48.1(tm544)*, *cdc-48.2(tm659)*, *brc-1(tm1145)*, and *brd-1(ok1623),* from the *Caenorhabditis* Genetics Center. We found that indeed the deficiency of each of these genes renders animals UV hypersensitive and causes increased developmental arrest upon UV, as shown by L1 larvae UV survival experiments (Fig. 5D-G, Supplementary Fig. 7). Next, we tested if their activity is epistatic to the TC-NER pathway or if the gene products act in a pathway parallel to TC-NER. Therefore, we crossed *wwp-1(ok1102), cdc48.1(tm544), brc-1(tm1145)*, and *brd-1(ok1625)* animals with *csb-1(ok2335)* animals. Interestingly, we found that each of the double mutants were more UV hypersensitive compared to their respective single mutants (Fig. 5D-G), as well as to the *csb-1(ok2335)* single mutant (Fig. 1A). These results suggest that WWP-1, CDC-48.1 and BRCA-1/BRD-1 function at least partially independent of TC-NER. We next crossed these mutants to the TC-NER deficient, but UV non-hypersensitive, *uvs-1(tm6134)* strain. Based on our FRAP experiments (Fig. 4D, E), we had concluded that this mutant is resistant to UV irradiation because Pol II is still effectively processed and removed from damaged DNA. Strikingly, we observed that the *uvs-1(tm6134)* double mutants with *wwp-1(ok1102), cdc48.1(tm544), brc-1(tm1145)* or *brd-1(ok1625)* were strongly hypersensitive to UV irradiation, to the same level as the *csb-1(ok2335)* double mutants. These results confirm that *uvs-1(tm6134)* animals are indeed TC-NER deficient but that their resistance to UV irradiation is due to Pol II processing involving the activity of WWP1, BRCA-1/BRD-1 and CDC-48.1.

## Discussion

Here, we use *C. elegans* to show that DNA damage in transcribed genes of non-dividing, somatic cells causes motoneuronal and developmental failure in TC-NER deficient animals due to the lack of proper Pol II processing and, consequently, the lack of DNA repair. These results are in line with previous observations by us and others showing that specifically TC-NER, but not GG-NER, protects against the severe consequences of DNA damage in somatic tissues, including muscles and neurons, and is therefore crucial to protect transcriptional integrity and normal animal development^67–69,73,75,84^. Interestingly, although TC-NER by itself is sufficient to protect developing animals against UV-induced DNA damage, the UV hypersensitivity of TC-NER mutants becomes more pronounced when in addition *xpc-1* is mutated (Fig. 1D), showing that GG-NER acts as backup in the repair of transcription-blocking DNA damage. Moreover, we found that GG-NER of transcription-blocking lesions in somatic tissues is very efficient but requires access to the DNA damage, which is prevented in most TC-NER deficient animals tested, due to impaired removal of Pol II from damaged DNA (Fig. 4G). We also show that this processing of lesion-stalled Pol II involves the activity of multiple E3 ubiquitin ligases, revealing that this is an intricate process that is tightly regulated at multiple levels (Fig. 5). Therefore, our data favor the idea that DNA damage-induced motoneuronal and developmental failure associated with TC-NER deficiency is primarily caused by the lack of lesion removal due to persistently stalled Pol II (Fig. 6), which may have important implications for understanding CS etiology and aging as discussed in more detail below.

**Fig. 6.**
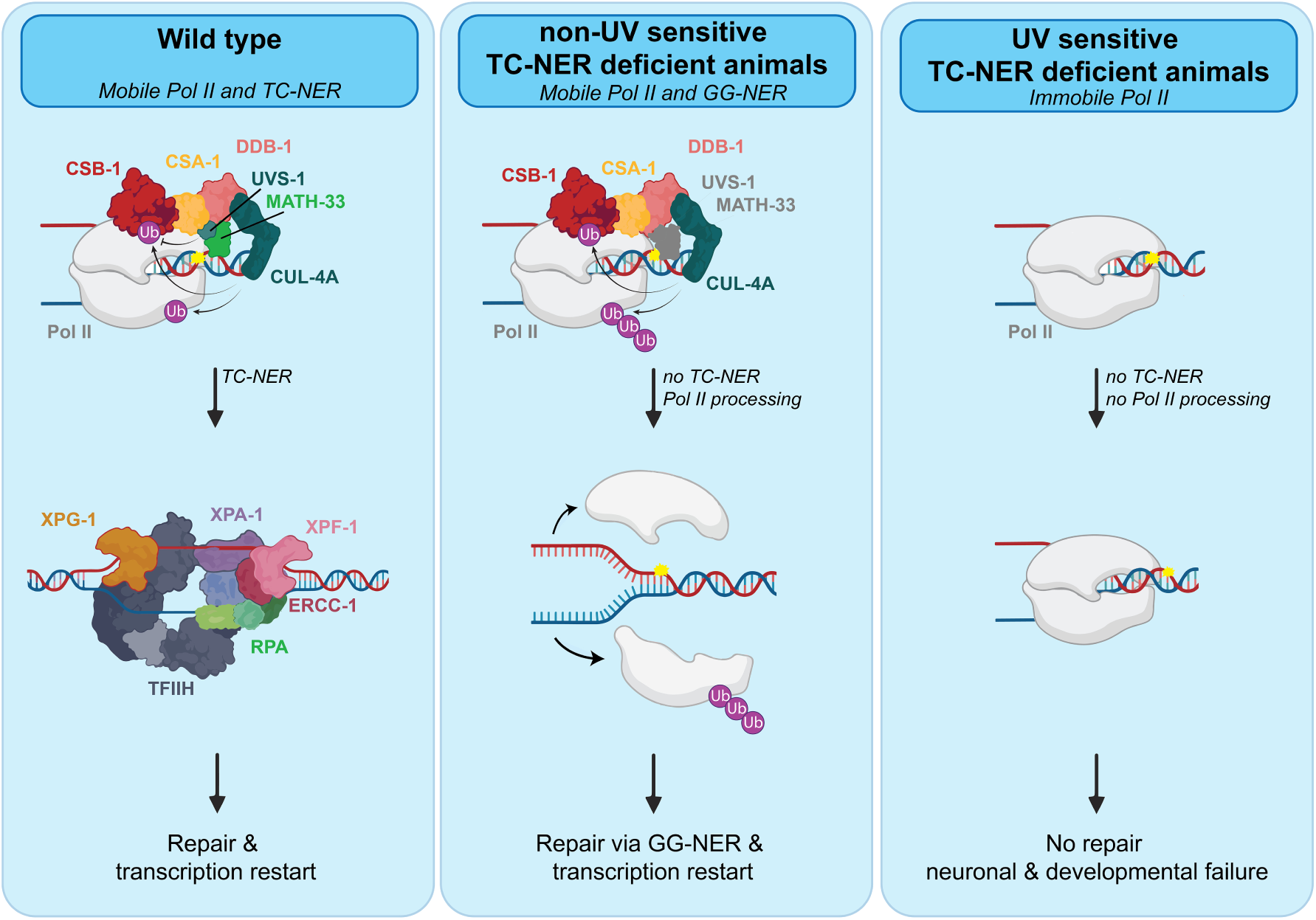
Model of Pol II stalling-induced transcription stress. Left panel: In wild type animals, lesion-stalled Pol II leads to the recruitment of the TC-NER machinery, Pol II is processed and subsequently the core NER machinery removes the lesion, facilitating transcription restart. Middle panel: Non-UV hypersensitive *uvs-1(tm6134)* mutant animals are TC-NER deficient, but Pol II is still processed, involving its ubiquitylation. This allows GG-NER to recognize and repair the transcription blocking lesion and subsequent transcription restart. Right panel: UV hypersensitive TC-NER deficient mutants, such as *csb-1* and *uvs-1* mutant animals, are also deficient in Pol II processing. This results in prolonged stalling of Pol II at DNA damage, shielding it from repair by other DNA repair pathways like GG-NER. The failure to process DNA damage-stalled Pol II leads to failure of transcription restart and, consequently, neuronal and developmental impairment in *C. elegans*. Ub stands for ubiquitin.

Transient depletion of persistently lesion-stalled Pol II rescued the UV-induced developmental arrest of TC-NER deficient animals in a GG-NER-dependent manner, confirming that GG-NER, via DNA damage detection by XPC-1, acts as backup to TC-NER in the removal transcription-blocking DNA damage. It is striking to note that *C. elegans* TC-NER and Pol II processing deficient mutants only exhibit a partial UV hypersensitivity compared to animals fully deficient in both TC-NER and GG-NER, such as *xpc-1; uvs-1* and *xpc-1; csb-1* double and *xpa-1* single mutants (Fig. 1D; 4G), and such as *gtf-2H5, xpg-1, ercc-1* and *xpf-1* mutants that we previously studied^67,68,84,91^. This suggests that even when Pol II is not properly processed, XPC-1 is still able to detect and initiate the removal of a subset of DNA lesions in transcribed genes. UV irradiation predominantly creates two types of DNA-helix-distorting lesions, i.e., cyclobutane-pyrimidine dimers (CPDs) and 6-4 pyrimidine-pyrimidone photoproducts (6-4PPs). 6-4PPs are rapidly removed by both TC-NER and GG-NER in mammalian cells, but CPDs are only rapidly removed by TC-NER and much slower by GG-NER, as XPC is unable to recognize these efficiently^111,112^. The partial UV sensitivity of TC-NER *C. elegans* mutants could therefore be due to inefficient CPD removal by GG-NER. This is, however, unlikely given the observation that transient Pol II depletion completely rescued the UV hypersensitivity of *csb-1(emc78)* and *uvs-1(emc80)* animals (Fig. 4G). These results suggest that, in *C. elegans,* XPC-1-mediated GG-NER is very efficient in repairing both 6-4PP and CPDs. Indeed, this is supported by our previous finding showing that CPDs are very rapidly cleared by GG-NER in *C. elegans* oocytes^67,68^. Moreover, it was previously shown that 6-4PPs and CPDs are repaired with equal efficiency in *C. elegans,* using radioimmunoassays on DNA extracted from whole animals^113^. Therefore, it is likely that in *C. elegans* somatic tissues, both TC-NER and GG-NER detect and repair UV-induced DNA damage, i.e. CPDs and 6-4PPs, in the template strand of active genes, and that it is a matter of chance whether XPC or Pol II first arrives at the lesion. In GG-NER deficient mutants, TC-NER is therefore sufficient to detect and remove all DNA damage in transcribed genes, explaining why GG-NER mutants are not UV hypersensitive in the L1 UV survival assay. However, in TC-NER deficient mutants, XPC-1 will be able to detect a fraction of the lesions in transcribed DNA, but Pol II will still be stalled by the remainder of lesions, shielding these from repair before XPC-1 has had the chance to detect these. This explains the partial UV hypersensitivity of TC-NER mutants and why these mutants become much more sensitive to UV irradiation in the absence of functional GG-NER. In view of this, it is interesting to note that single TC-NER deficient *Csb^−/−^* and *Csa^−/−^* mice display only mild CS features, but that additional inactivation of GG-NER by either *Xpc* or *Xpa* inactivation dramatically aggravates their CS phenotype, including neurological degeneration, motor dysfunction, and progeroid features^114–116^. This has been attributed to increased DNA damage load due to additional XPC or XPA mutation, which in essence shows that also in mice GG-NER acts as backup pathway to TC-NER. Taken together, these results demonstrate that both subpathways of NER, GG- and TC-NER, act in synergy to repair DNA damage in actively transcribed genes in somatic, nondividing tissues such as neurons.

Remarkably, we found that TC-NER deficiency does not necessarily lead to developmental arrest after UV irradiation, as observed in the *uvs-1(tm6134)* animals. These mutant animals are clearly TC-NER deficient, as evidenced by their resistance to trabectedin (Fig. 1G), but showed a UV sensitivity (Fig. 1C) and UV-dependent Pol II mobility (Fig. 4D-E) comparable to wild type animals. In contrast to wild type animals, however, their UV sensitivity was fully dependent on *xpc-1* (Fig. 1D). This indicates that in these mutants, Pol II does not persistently stall at transcription-blocking lesions and that XPC-1 has full access to initiate repair via GG-NER (Fig. 6). It is therefore likely that the *tm6134* mutant UVS-1 protein, together with USP7, is still recruited to the TC-NER complex, despite the fact that in this mutant the VHS domain, which is necessary for interaction with CSA^30^, is partially deleted. Indeed, in human cells also CSA-independent recruitment of UVSSA to DNA damage has been observed^80^. In contrast, the TC-NER deficient *uvs-1(tm6311)* mutant animals displayed strong motoneuronal and developmental failure upon UV irradiation. As in these mutants specifically the TRAF-binding motif and most of the p62/GTF-2H1-binding region are deleted, their UV hypersensitivity could be caused by the inability to recruit the deubiquitylase enzyme USP7 and/or TFIIH to the TC-NER machinery. In mammalian cells, loss of USP7 recruitment to the TC-NER machinery results in increased CSB degradation, as consequence of its ubiquitylation by the CRL4^CSA^ complex^27,28,32,37,92^. In line with this, we found that CSB-1 protein levels were greatly reduced in *C. elegans uvs-1(tm6311)* oocytes, but not in *uvs-1(tm6134)* animals (Fig. 3D-E). Still, overexpression of CSB-1 did not rescue the UV-induced developmental arrest of *uvs-1(tm6311)* mutants, suggesting that also TFIIH recruitment is important. Indeed, animals with a mutation in the p62/GTF-2H1-binding region only were as UV hypersensitive as animals with a mutation in the TRAF-binding motif only. The recruitment of CRL4^CSA^ by CSB^9,10^ as well as the helicase/translocase activity of TFIIH^4,93–98^ are likely needed for the processing of stalled Pol II from the lesion, suggesting that the UV-hypersensitivity in *uvs-1* mutant animals results from the inability to efficiently process Pol II. Indeed, prolonged Pol II immobilization was observed upon UV-irradiation in *uvs-1(tm6311)* animals (Fig. 4D-E).

As our results indicate that not TC-NER deficiency perse, but rather DNA damage-induced persistent Pol II stalling causes developmental arrest, it is evident that processing of lesion-stalled Pol II must be tightly regulated. Indeed, to safeguard transcriptional integrity, several E3 ubiquitin ligases have been implicated in ubiquitylating and regulating Pol II processing^33,59–63,104–106^, but their precise function and interplay are still unclear. Here, we show that the loss of subunits of the E3 ubiquitin ligases CRL4^CSA^, NEDD4, CRL5^Elongin^, CRL2^VHL^, and BRCA1/BARD1, as well as of the ubiquitin-dependent segregase VCP, all results in UV-induced developmental arrest in *C. elegans,* likely in a CSB-dependent^104^, but also CSB-independent manner. Removal of lesion-stalled Pol II by polyubiquitylation and degradation, involving the sequential activities of NEDD4 and CRL5^Elongin^, has been proposed to act as an alternative last resort pathway, at least in yeast, to failing TC-NER^57,59,60,117^. However, our observation that RNAi knockdown or mutation of single subunits of these E3 ubiquitin ligases already causes UV hypersensitivity suggests that their activity in Pol II processing is relevant for TC-NER itself as well. Our findings indicate that multiple different E3 ubiquitin ligases, together with CRL4^CSA^, coordinate Pol II displacement and/or degradation to enable lesion removal. Further research is needed to mechanistically understand the individual roles of each of these E3 ubiquitin ligases in Pol remodeling and TC-NER, their interplay and how their collective activity is controlled. Also, it will be necessary to study to which extent they act in a CSB-dependent or CSB-independent manner.

Hereditary defects in human CSB and CSA can lead to severe CS, while hereditary mutations in human UVSSA lead to the much milder UV^s^S. We and others have previously put forward the hypothesis that CS features are therefore not caused by mere TC-NER deficiency, but by the failure to properly remove stalled Pol II from DNA damage^4,30,31,33,52–54^. Indeed, it was shown that CSA and CSB deficient cells differ from UVSSA deficient cells in their ability to remove Pol II from chromatin after DNA damage induction^55,56,118^. Our results in this manuscript strongly support this model by directly showing that Pol II processing is necessary to facilitate DNA repair and to prevent DNA damage-induced neuronal and developmental failure. We therefore propose that severe CS features in humans are, at least partially, due to persistently stalled Pol II, which shields DNA damage from repair by alternative means and causes transcriptional failure (Fig. 6). This hypothesis implies that CS features are ultimately caused by a failure to repair DNA damage, albeit not only a failure of TC-NER itself but also of other repair mechanisms. Defects in downstream NER factors XPB, XPD, XPF and XPG can also cause CS features, as observed in patients with Xeroderma pigmentosum-Cockayne syndrome (XP-CS) complex^38,51,119,120^. We recently showed that TFIIH persistently binds to DNA damage in XPF- and XPG-deficient cells, which leads to more severe cellular defects after DNA damage induction^91^. This suggests that possibly the accumulation of persistently stalled NER intermediates, preventing repair by other means, is a common underlying cause of CS features. The progeroid nature of these features, which resemble genuine human aging, is one of the reasons why accumulating DNA damage is considered a central driver of aging^6,121^. Especially neurons are affected by DNA damage in both in CS and in aging. This is likely because of their postmitotic status, long lifespan and the fact that they express some of the longest genes, which makes them especially susceptible to the accumulation of DNA damage interfering with transcription ^7,122,123^. It is not entirely clear why DNA damage accumulates with age, and to what extent declining DNA repair rates are responsible for this^124^. Interestingly, it was found that a high percentage of Pol II complexes is stalled at DNA damage in aging mice, which depends on gene length and leads to gene expression changes associated with aging^7^. Possibly, this may point to the fact that not only DNA repair itself, but also Pol II processing capacity declines with age, which may lead to more Pol II stalling and stronger inhibition of repair, causing persistent DNA damage-induced transcription stress that leads to aging. For this reason, it will be interesting to study mechanisms and consequences associated with Pol II processing during aging as well.

## Materials and methods

### *C. elegans* strains and culture

All *C. elegans* strains were cultured according to standard methods^125^ on nematode growth media (NGM) agar plates seeded with *Escherichia coli* OP50. Bristol N2 was used as wild type strain. All strains used are listed in Supplementary Table 1. All mutants generated were backcrossed against wild type and genotyped by PCR and sequencing. Double mutants were generated by crossing and checked by genotypic PCR. During all assays, animals were kept at 20°C. Mutant animals were generated by injection of a mixture of Cas9 and gRNA (Integrated DNA technologies (IDT)) and, in case of knock-in and point mutants, a homology directed repair template, or, in case over CSB overexpression, a mixture of the *csb-1::FLAG::GFP* PCR product and fluorescent marker *elt-2::mCherry*. All generated knock-out, knock-in, point mutant and transgenic animals were verified by genotyping PCR and sequencing. *csb-1(emc78)* and *uvs-1(emc80)* mutants, in which the complete *csb-1* or *uvs-1* genes were deleted, respectively, were generated using sgRNAs targeting *csb-1* (5’-TGAAAAAATACCTAAGTACC-3’ and 5’-AAAAATGAATCAATGAATAA-3’) or *uvs-1* (5’-CAAATAAAATGTTGAAAAGA-3’ and 5’-CAGTTTTCTCATTTTTAATA-3’). To generate knock-in animals, wild type animals, unless mentioned otherwise, were injected with gRNA, a homology directed repair template consisting of a heteroduplex DNA fragment that was generated by mixing PCR products^126^, and purified Cas9 protein (IDT). To generate *uvs-1::AID::GFP* knock-in wild type, *uvs-1(tm6134)*, and *uvs-1(tm6311)* animals, a gRNA with targeting sequence 5’-CAGTTTTCTCATTTTTAATA-3’ (IDT) was used. The homology directed repair template was generated using primer combinations 5ʹ-CGTCTAGAAAAAAATTTTGGTCAACAGTTTTCTCATTTTATGCCTAAAGATCCAGCCAAAC-3ʹ/5ʹ-ATATTTACTTTATTTTTCTAAATTTTAACAAAAAACCGTATCATTTGTATAGTTCGTCC-3ʹ and 5ʹ-ATGCCTAAAGATCCAGCCAA-3ʹ/5ʹ-TCATTTGTATAGTTCGTCCATGCC-3ʹ on a gene fragment consisting of *AID::GFP* sequences flanked by 37 bp left and 33 bp right homology arms from the *uvs-1* locus (generated by IDT). To generate *csb-1::AID::GFP* and *csb-1::AID::mScarlet3* knock-in animals, a gRNA with targeting sequence 5’-TTTTACATTCTAGAAGTACT-3’ (IDT) was used. The homology directed repair of AID::GFP was generated using primer combinations 5ʹ-CGGCAGATGTGCTGTCGCCAG-3ʹ/5ʹ-GCAATTTGGGGCTGGGGGATTC-3ʹ and 5ʹ-ATGCCTAAAGATCCAGCCAA-3ʹ /5ʹ-TCATTTGTATAGTTCGTCCATGCC-3ʹ on a gene fragment consisting of *AID::GFP* sequences flanked by 110 bp left and 93 bp right homology arms from the *csb-1* locus (generated by IDT). The homology directed repair of AID::mScarlet3 was generated using primer combinations 5’-GGTCGGATTCATCAGGACCGAG-3’/5’-GCTCCCATTATTCATTGATTCATTTTTCAC-3’ on a gene fragment consisting of AID::mScarlet sequences flanked by 35 bp left and 80 bp right homology arms from the *csb-1* locus (generated by IDT). *AID::3xFLAG::GFP::ama-1* knock-in animals were generated by injecting in previously generated *ama-1(ot1037[GFP::3xFLAG::::ama-1])*^127^ animals, using gRNA 5’ TTAAATTTTCAGATGAGTAA 3’ and a template containing AID sequences generated by PCR with primers 5’-CAATTTTTTTAAATCCCATTTTTTAAATTTTCAGATGCCTAAAGATCCAGCCAAAC-3’ and 5’-GACAACTCCAGTGAACAATTCTTCTCCTTTACTCATCTTCACGAACGCCGCCGCCTC-3’. *ama-1*(*em216*) animals were generated using a gRNA with targeting sequence 5’-CGAATCGCCGGAGAGGATAA-3’ and as homology directed repair template ssDNA oligomers with sequence GATCGGATTCACGGAGGCTTCGGCAACGATGTTCACACTATCTACACCGACGATAACGCCGAGAAGCTCGTTTTCCGTCTCAGGATAGCGGGGGAAGACAGAGGGGAAGCTCAGGAGGAGCAGGTGGATAAGATGGAGGACGACGTGTTCCTACGATGTATCGAGGCAAATATGTTGTCAGATTTGACGCTTCAGGGAATC. Animals overexpressing CSB-1::GFP were generated by injecting a fusion PCR product, combining genomic *csb-1* sequence, obtained by PCR using primer combinations AGTGACTGTTGACGAATCCAGC and TCGTCGTCATCCTTGTAGTCCATTCTAGAAGTACTCGGTCCTGATGAA, with FLAG-GFP sequences. Following injection, the extrachromosomal transgene was integrated using ionizing radiation^128^. For overexpression of UVS-1::GFP, a PCR product including the endogenous promoter (1901 bp) was generated from genomic DNA of each respective *uvs-1::AG* knock-in animals, using primers CTTGGACTCATTTAGCTGCG and CGGGTGCGGGTGGACAACCTC (wild type: 5520bp, *tm6134:* 5079 bp, *tm6311:* 4997 bp), and injected into the full knock out *uvs-1(emc80)* mutant.

### RT-PCR and RT-qPCR

For RT-PCR, animals were lysed in TRIzol (Qiagen) and RNA was purified using RNeasy spin columns (Qiagen). cDNA was generated using Superscript II Reverse Transcriptase (Invitrogen) according to manufacturer’s instructions. For RT-qPCR, RNA was isolated and cDNA generated from ten young adult animals per strain and condition using the Power SYBR Green Cells-to-Ct kit (Invitrogen) according to previous described method^129^. qPCR was performed using PowerUp SYBR Green Master Mix (Invitrogen), according to manufacturer’s instructions, with 58 °C as annealing temperature and 1 min elongation time. Primers used for RT-PCR and RT-qPCR are listed in Supplementary Table 2. cdc-42 and pmp-3 were used as reference genes.

### L1 larvae UV survival assay

L1 larvae UV survival experiments were performed as previously described^67,71^. In brief, eggs were collected from adult animals by hypochlorite treatment and plated on five 6 cm NGM plates, seeded with HT115(DE3) bacteria, per UV dose. After 16 h, L1 larvae were irradiated at the indicated UV-B doses (Philips TL-12 tubes, 40W). Following a 48 h recovery period, the number of arrested (L1/L2 larvae) and developed (L3/L4/young adult) animals were counted and survival percentage was calculated. For depletion with the auxin derivative 5-Ph-IAA, L1 larvae were irradiated with 60 J/m^2^ UV-B. After 4 h recovery, the irradiated L1 larvae were plated for 4 h on 6 cm NGM plates seeded with HT155(DE3) and containing 50 µM 5-Ph-IAA. Next, the animals were transferred to 6 cm NGM plates seeded with HT115(DE3) bacteria. After 40 h recovery, the number of arrested (L1/L2 larvae) and developed (L3/L4/young adult) animals were counted and survival percentage was calculated.

### Trabectedin survival assay

Trabectedin (MCE) survival experiments were performed by collecting eggs from adult animals by hypochlorite treatment and hatching these in M9 buffer containing the indicated trabectedin concentrations, while rotating at room temperature for 16 h. After hatching, L1 larvae were washed twice with M9 buffer, and divided over three 6 cm NGM plates, seeded with HT115(DE3) bacteria, per concentration. After a 48 h recovery period, the number of arrested (L1/L2 larvae) and developed (L3/L4/young adult) animals was counted and survival percentages were calculated.

### Motility assay

To assess neuromotor function, staged first day adult animals were mock treated or UV-irradiated (60 J/m^2^ UV-B) and cultured for 72 h (Fig. 1H), after which the animals were placed in 5 μl M9 buffer and allowed to acclimatize for 30 s before the number of lateral body movements was counted for 30 s.

### Real-time imaging and FRAP in *C. elegans*

For microscopy and fluorescence measurements of GFP and mScarlet3 knock-in strains, staged living animals were mounted on 2% agar pads in M9 buffer containing 10 mM NaN_3_ (Sigma) and imaged on a Leica TCS SP8 microscope (Leica Microsystems). For measurement of CSB-1::AID::mScarlet3 fluorescence levels, animals were washed three times with M9 buffer and mock treated or irradiated (300 J/m^2^ UV-B), after which animals were allowed to recover on NGM plates seeded with OP50 bacteria for 2 h. Fluorescence levels were quantified by determining the mean mScarlet3 intensity minus background levels of individual oocyte nuclei using ImageJ software. CSB-1 levels in the four most proximal oocytes were quantified. For measurement of AGF::AMA-1 mobility by FRAP, staged adult animals were immobilized, using a mixture of M9 buffer, levamisole (10 mM; Sigma) and polystyrene beads (Polyscience Inc.) on 6% agarose pads. For UV irradiation, animals were first washed three times with M9 buffer and irradiated on empty agar plates (80 J/m^2^ UV-B) and allowed to recover on NGM plates seeded with OP50 bacteria for the indicated time periods. Animals were imaged using a Leica TCS SP8 microscope (Leica Miscrosystems) for a period of maximum 30 min. For FRAP, intestinal and hypodermal nuclei were imaged at 1400 Hz using a 488 nm laser at low power for 1.88 s. Next, GFP fluorescence in a small square (1.0 × 1.0 μm, zoom 10) was photobleached using high laser power. The photobleaching in the small square was first optimized such that the reduction in the overall nuclear fluorescence signal was only minimal.

Fluorescence recovery was measured at low laser power every 18 ms until steady-state levels were reached. Fluorescent signals were normalized to the average fluorescence intensity before photobleaching and plotted as average of at least 2 to 3 cells per animal, for a total of at least 9 animals per condition, obtained in at least three independent experiments. The immobile fractions (F_imm_) were determined using the fluorescence intensity measured immediately after UV (I_0_) and the average steady-state fluorescent signal after complete recovery, from untreated (I_final, unt_) and UV-treated cells (I_final, UV_), and applying the formula:

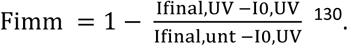

LAS AF software (Leica) was used for imaging and ImageJ software for quantification.

### RNAi experiments

For RNAi knockdown experiments, RNAi bacteria from the *Caenorhabditis* elegans RNAi feeding library^110^ were used. Control RNAi was vector pPD129.36 (a gift from Andrew Fire). RNAi knockdown was accomplished by growing wild type animals on NGM plates containing HT115(DE3) bacteria expressing gene-specific RNA for three generations before survival experiments were performed. Survival experiments were as described for L1 larvae UV survival assay.

### Protein prediction and alignments

Protein prediction were obtained from FGENESH ^83^ and corresponding protein alignments were generated through CLUSTAL O (1.2.4) multiple sequence alignment and edited in Jalview 2.11.3.0.

### Quantification and Statistical Analysis

Mean values and error bars indicating S.E.M. are shown for each experiment. Statistical significance was determined by two-way ANOVA followed by Šídák’s multiple comparisons test, as indicated in the legends for each experiment. All analyses were performed using Graph Pad Prism version 9 for Windows (GraphPad Software, La Jolla California USA).

## Acknowledgements

We thank Dr. G. Jansen for use of his injection microscope, Dr. Andrew Fire for plasmids and the Erasmus MC Optical Imaging Center for microscope support. Some strains were provided by the Caenorhabditis Genetics Center (funded by NIH Office of Research Infrastructure Programs P40 OD010440) and the National Bioresource Project for the nematode. This work was supported by the Netherlands Organization for Scientific Research (711.018.007).

## Author contributions

MvdW and KLT performed all experiments. MS performed experiments that initiated this study. MvdW, KLT and HL designed experiments. JAM, WVM and HL conceptualized ideas and supervised experiments. MvdW and HL wrote the manuscript. All authors reviewed the manuscript.

## Competing interests

The authors declare no competing interests

## Notes

### Competing Interest Statement

The authors have declared no competing interest.

