## Supplemental information for "RNA polymerase II processing facilitates DNA repair and prevents DNA damage-induced neuronal and developmental failure"

**A**

| | |
|---|---|
| CSB-1 | MSTKPE………GPSTSRM* |
| CSB-1(emc78) | MKKPELTATSVEKFLIEKF* |

**B**

| | |
|---|---|
| UVS-1 | MLKRRQPK…..QFSHF* |
| UVS-1(emc80) | MKKPELTASTRPRAKE* |

**Supplementary Figure 1.** Predicted protein sequences encoded by full knock-out alleles of (**A**) *csb-1(emc78)* and (**B**) *uvs-1(emc80)*, as compared to the N- and C-terminal protein sequences encoded by the wild type CSB-1 and UVS-1 alleles. Red color indicates nonsense sequence. Black asterisks indicate wild-type stop codon at position 582 for UVS-1 and 957 for CSB-1, red asterisks indicate stops codons for both mutants.

```
                               VHS domain
UVS-1             MDSIDNSTTIIRKNLNRFIRELTDDGKLDFESIPYQNLQKEVANQDEEGCENVIEVLLDT  60
UVS-1 (tm6134)    MDSIDNSTTIIRKNLNRFIRELTDDGKLDFESIPYQNLQKEVANQDEEGCENVIEVLLDT  60
UVS-1 (tm6311)    MDSIDNSTTIIRKNLNRFIRELTDDGKLDFESIPYQNLQKEVANQDEEGCENVIEVLLDT  60

                               VHS domain
UVS-1             TSRSGCPDRKLILQLFNSFFLQFPIFRENLLNDPSEFLELMFETNPIRNPLPGSKKHGNE  120
UVS-1 (tm6134)    TS----------------------------------------------------------  62
UVS-1 (tm6311)    TSRSGCPDRKLILQLFNSFFLQFPIFRENLLNDPSEFLELMFETNPIRNPLPGSKKHGNE  120

                       VHS domain
UVS-1             LKVEAITVIKSWEKEKCVKNDARMKCLVVTLKKTKFVDYENGAKKIEAERKRKKILEERK  180
UVS-1 (tm6134)    ------------------------------------------------------------  62
UVS-1 (tm6311)    LKVEAITVIKSWEKEKCVKNDARMKCLVVTLKKTKFVDYENGAKKIEAERKRKKILEERK  180

UVS-1             MKMIENSVNVYSSKYHEIKNDAETLSMELTTTMQMLVPSFTTADPEVPSTSTSTPSAISD  240
UVS-1 (tm6134)    ------SVNVYSSKYHEIKNDAETLSMELTTTMQMLVPSFTTADPEVPSTSTSTPSAISD  116
UVS-1 (tm6311)    MKMIENSVNVYSSKYHEIY-----------------------------------------  199

UVS-1             SKSFEIFIPDLTPEISVSSENDAIVEAFLGAKLSLIHRVQTLRKLVKRLQLLKQPGEKLA  300
UVS-1 (tm6134)    SKSFEIFIPDLTPEISVSSENDAIVEAFLGAKLSLIHRVQTLRKLVKRLQLLKQPGEKLA  176
UVS-1 (tm6311)    ------------------------------------------------------------  199

UVS-1             QEIIDYRDGIKNLVLKADELRIINPRPPKNKRKKSDDDFIDVDISIDDILMVQYAEKLEV  360
UVS-1 (tm6134)    QEIIDYRDGIKNLVLKADELRIINPRPPKNKRKKSDDDFIDVDISIDDILMVQYAEKLEV  236
UVS-1 (tm6311)    --------------------------------------------SIDDILMVQYAEKLEV  215

                                                   DUF2043 domain
UVS-1             DVKSKDESEKITESPEKHKIEMKNEKPVKIKTVPFGLDLKYWGEERKDVEVPKNNADCHR  420
UVS-1 (tm6134)    DVKSKDESEKITESPEKHKIEMKNEKPVKIKTVPFGLDLKYWGEERKDVEVPKNNADCHR  296
UVS-1 (tm6311)    DVKSKDESEKITESPEKHKIEMKNEKPVKIKTVPFGLDLKYWGEERKDVEVPKNNADCHR  275

                               DUF2043 domain
UVS-1             FWRSADEGTVAGKAQQSIYTQRQYTFIGKAPDNRKVCLAKMKSGKLCPRKDYYTCPLHGK  480
UVS-1 (tm6134)    FWRSADEGTVAGKAQQSIYTQRQYTFIGKAPDNRKVCLAKMKSGKLCPRKDYYTCPLHGK  356
UVS-1 (tm6311)    FWRSADEGTVAGKAQQSIYTQRQYTFIGKAPDNRKVCLAKMKSGKLCPRKDYYTCPLHGK  335

                   DUF2043 domain
UVS-1             IVDRDDEGRPINEEDRLEENYRKEQNHLKEADKIRQMIEKEYESKTKRRKKHDVDTTASE  540
UVS-1 (tm6134)    IVDRDDEGRPINEEDRLEENYRKEQNHLKEADKIRQMIEKEYESKTKRRKKHDVDTTASE  416
UVS-1 (tm6311)    IVDRDDEGRPINEEDRLEENYRKEQNHLKEADKIRQMIEKEYESKTKRRKKHDVDTTASE  395

UVS-1             DVRNRLQKKLLDPKTIQRVSADLDASRKNRLEKNFGQQFSHF                    582
UVS-1 (tm6134)    DVRNRLQKKLLDPKTIQRVSADLDASRKNRLEKNFGQQFSHF                    458
UVS-1 (tm6311)    DVRNRLQKKLLDPKTIQRVSADLDASRKNRLEKNFGQQFSHF                    437
```

TRAF-binding motif (Higa et al., 2016)

p62/GTF-2H1-binding domain (Okuda et al., 2017)

**Supplementary Figure 2.** Predicted protein sequences with f functional domains of wild type UVS-1 compared to mutant proteins encoded by the *uvs-1(tm6134)* and *uvs-1(tm6311)* alleles. The TRAF-binding motif, essential for USP7 binding, is indicated in red and the p62/GTF-2H1-binding region in blue.

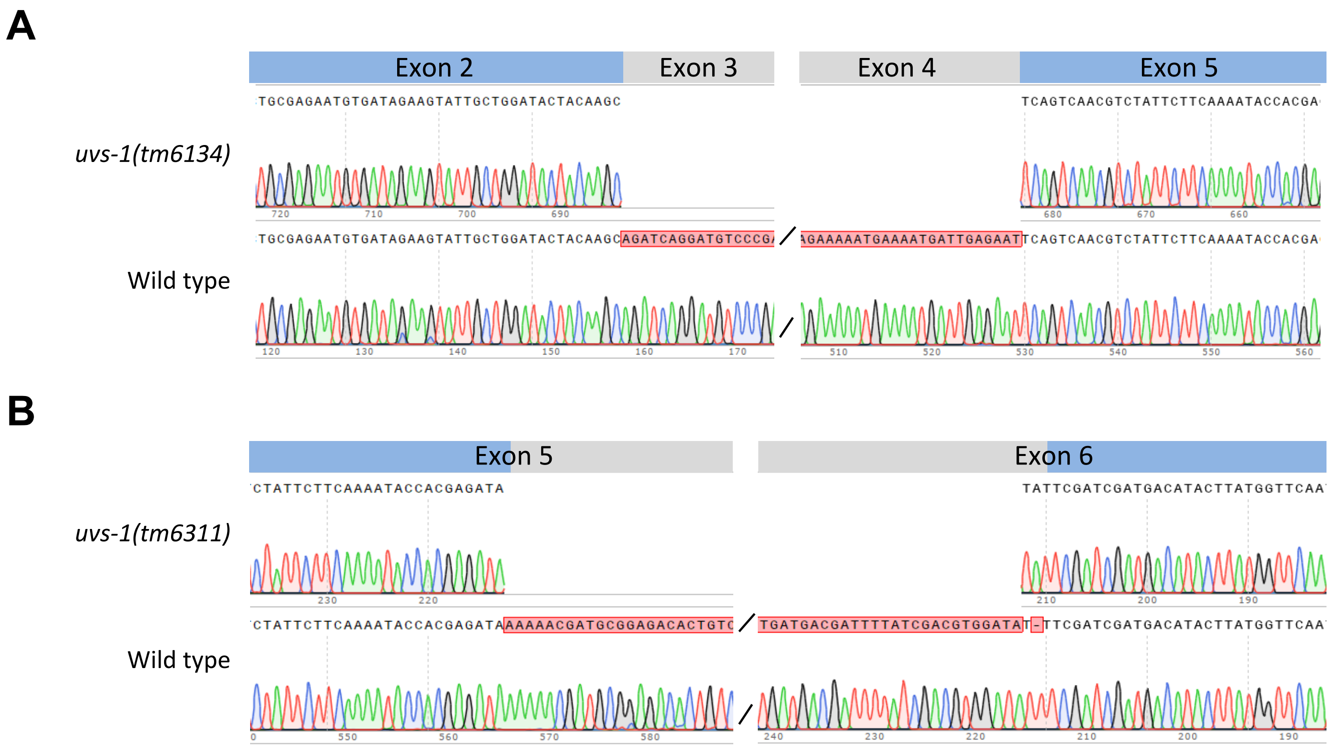


**Supplementary Figure 3.** cDNA sequencing of (**A**) the *uvs-1(tm6134)* allele shows that this allele leads to an in-frame deletion of exon 3 and 4 in which exon 2 is connected to exon 5 and of (**B**) the *uvs-1(tm6311)* allele shows that this allele leads to an in-frame partial deletion of exons 5 and 6 with an insertion of two nucleotides.

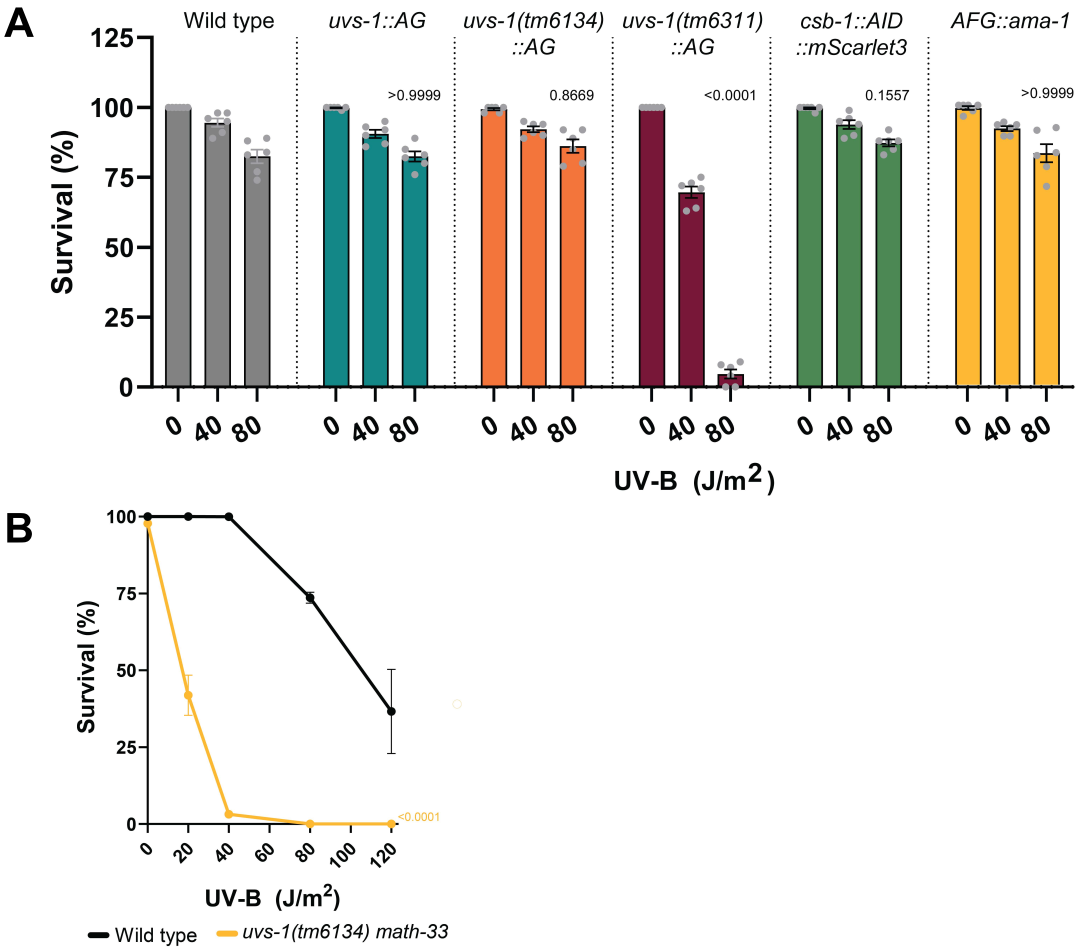


**Supplementary Figure 4.** L1 larvae UV survival assays of **(A)** wild type and AID::GFP (AG) or AID::GFP:: 3xFLAG (AGF) expressing knock-in animals, including *uvs-1::AG, uvs-1(tm6134)::AG, uvs-1(tm6311)::AG, csb-1::AID::mScarlet3,* and *AGF::ama-1;* and of (**B**) wild type and *uvs-1(tm6134) math-33* animals. Mean with SEM of three independent experiments. Numbers in the graph represent *p-values* determined by TWO-WAY ANOVA Šídák's multiple comparisons test.

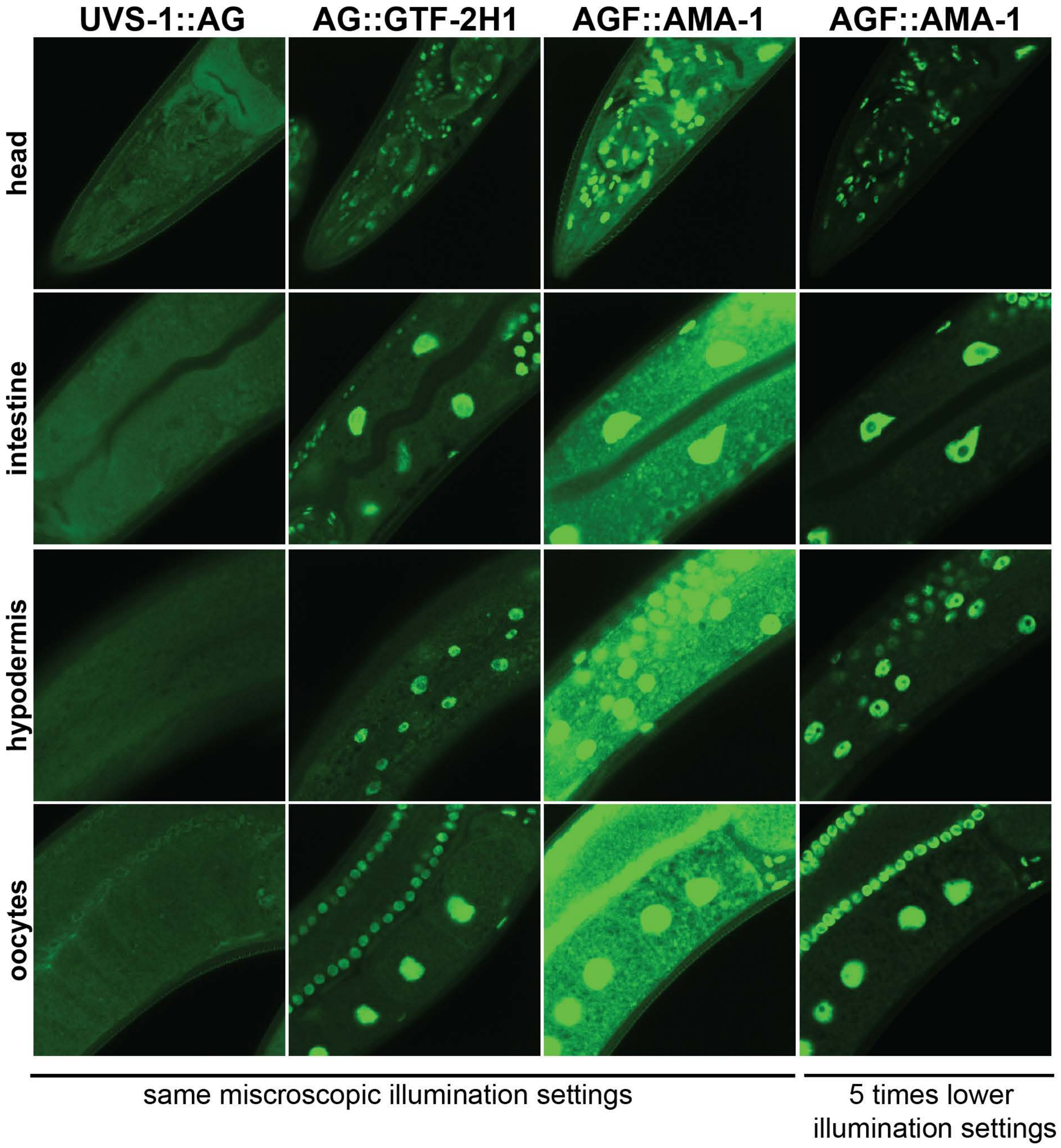


**Supplementary Figure 5.** Representative fluorescence confocal microscopy pictures, imaged using the same confocal microscopy settings (first three rows) or after lowering the intensity five times (fourth row), of living knock-in animals expressing UVS-1::AG, AG::GTF-2H1, or AGF::AMA-1 in the head, intestine, hypodermis, oocytes, and embryos.

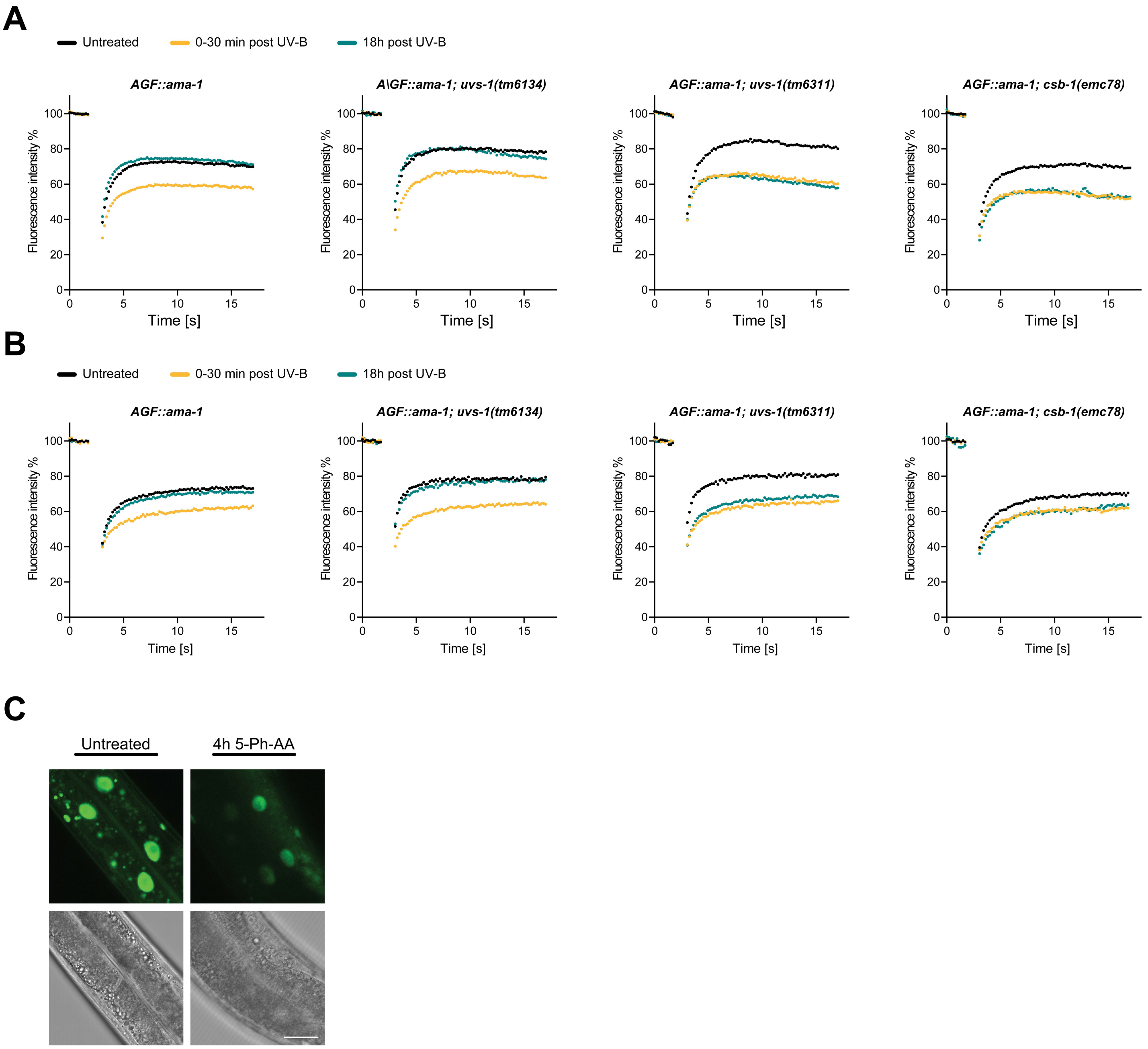


**Supplementary Figure 6.** Fluorescence recovery after photo-bleaching (FRAP) experiments in (**A**) hypodermal and (**B**) intestinal nuclei of living AGF::AMA-1-expressing animals, either untreated (black) or immediately (0-30 min; yellow) or 18 h (blue) after induction of DNA damage by UV irradiation. FRAP was measured in wild type, *uvs-1(tm6134)*, *uvs-1(tm6311)*, and *csb-1(emc78)* genetic backgrounds. Immobile fractions calculated based on these curves are shown in Figure 4D-E. Each line shows the mean of measured cells in three independent experiments for untreated, 0-30 min post UV-B, and 18 h post UV for (**A**) WT (n=51, 37, 47), *uvs-1(tm6134)* (n=22, 22, 23), *uvs-1(tm6311)* (n=22, 20, 25), and *csb-1(emc78)* (n=23, 21, 22), for (**B**) WT (n=47, 35, 45), *uvs-1(tm6134)* (n=20, 33, 21), *uvs-1(tm6311)* (n=21, 31, 29), and *csb-1(emc78)* (n=27, 32, 20). (**C**) Representative fluorescence confocal microscopy images of AGF::AMA-1 expression in intestinal nuclei of adult knock-in animals that were untreated or treated for 4 h with 5-Ph-IAA. Scale bar: 50 µm.

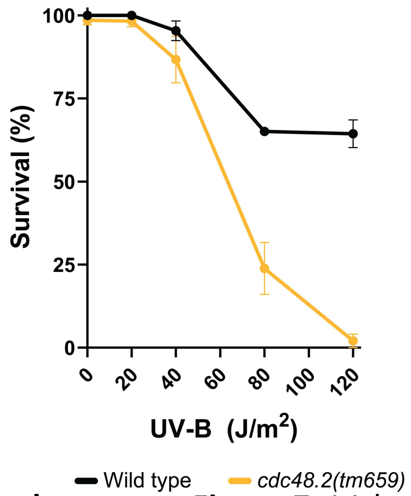


**Supplementary Figure 7.** L1 larvae UV survival assay of wild type and *cdc-48.2(tm659)* animals. The percentage survival of animals that developed beyond the L2 larvae stage are plotted against the applied UV-B doses. Mean with SEM of three independent experiments. Numbers in the graph represent *p-values* determined by TWO-WAY ANOVA Šídák's multiple comparisons test.

**Supplementary Table 1** ***C. elegans strains***

| Strain | Genotype |
|---|---|
| **Bristol N2** | wild type |
| **DW102** | brc-1(tm1145) III |
| **FX544** | cdc-48.1(tm544) II |
| **FX659** | cdc-48.2(tm659) II |
| **FX2026** | polq-1(tm2026) III |
| **FX5232** | csa-1(tm5232) II |
| **GJ1534** | xpc-1(tm3886) IV; csb-1(ok2335) X |
| **GJ1553** | xpc-1(tm3886) IV |
| **HAL25** | uvs-1(tm6134) V |
| **HAL26** | csb-1(ok2335) X |
| **HAL34** | wwp-1(ok1102) I; uvs-1(tm6134) V |
| **HAL113** | xpc-1(tm3886) IV; uvs-1(tm6134) V |
| **HAL120** | wwp-1(ok1102) I; csb-1(ok2335) X |
| **HAL121** | uvs-1(tm6311) V |
| **HAL255** | csb-1(emc78) X |
| **HAL257** | uvs-1(emc80) V |
| **HAL275** | cshIs140 [P(rps-28)::TIR1[F79G]::mCherry] II; ama-1(emc86[AID::GFP::ama-1]) [ot1037] IV |
| **HAL278** | cshIs140 [P(rps-28)::TIR1[F79G]::mCherry] II; ama-1(emc86[AID::GFP::ama-1]) [ot1037] IV; csb-1(emc78) X |
| **HAL279** | cshIs140 [P(rps-28)::TIR1[F79G]::mCherry] II; ama-1(emc86[AID::GFP::ama-1]) [ot1037] I; uvs-1(emc80) V |
| **HAL280** | xpa(ok698) I; cshIs140 [P(rps-28)::TIR1[F79G]::mCherry] II; ama-1(emc86[AID::GFP::ama-1]) [ot1037] IV |
| **HAL293** | ama-1(emc86[AID::GFP::ama-1]) [ot1037]) IV; csb-1(emc94[HA::csb-1]) X |
| **HAL294** | uvs-1(emc95[uvs-1::AID::GFP]) V |
| **HAL296** | ama-1(emc86[AID::GFP::ama-1]) [ot1037]) IV; csb-1(emc96[csb-1::HA]) X |
| **HAL298** | polq-1(tm2026) III; uvs-1(tm6134)V |
| **HAL300** | polq-1(tm2026) III; uvs-1(emc97[uvs-1(tm6134)::AID::GFP]) V |
| **HAL260** | gtf-2H1(emc202[AID::GFP::gtf-2H1]) IV |
| **HAL505** | elof-1(emc203) II |
| **HAL514** | brc-1(tm1145) III; uvs-1(tm6134) V |
| **HAL515** | cdc-48.1(tm544) II; uvs-1(tm6134) V |
| **HAL516** | brd-1(ok1623) III; uvs-1(tm6134) V |
| **HAL519** | brc-1(tm1145) III; csb-1(ok2335) X |
| **HAL520** | brd-1(ok1623) III; csb-1(ok2335) X |
| **HAL521** | cdc-48.1(tm544) II; csb-1(ok2335) X |
| **HAL543** | ama-1(emc216[K1260R]) IV |
| **HAL546** | ama-1(emc86[AID::GFP::ama-1]) [ot1037] IV; uvs-1(tm6311) V |
| **HAL547** | ama-1(emc86[AID::GFP::ama-1]) [ot1037] IV; uvs-1(tm6134) V |
| **HAL548** | uvs-1(emc214[uvs-1(tm6311)::AID::GFP]) V |
| **HAL549** | polq-1(tm2026) III; csb-1(emc215[csb-1::AID::mScarlet3]) X |
| **HAL550** | csb-1(emc215[csb-1::AID:mScarlet3) X |
| **HAL551** | cshIs140[P(rps-28)::TIR1[F79G]::mCherry] II; xpc-1(tm3886) ama-1(emc86[AID::GFP::ama-1]) [ot1037] IV; csb-1(emc78) X |
| **HAL552** | cshIs140[P(rps-28)::TIR1[F79G]::mCherry] II; xpc-1(tm3886) ama-1(emc86[AID::GFP::ama-1]) [ot1037] IV |
| **HAL554** | uvs-1(tm6134) V; csb-1(emc215[csb-1::AID:mScarlet3]) X |
| **HAL555** | uvs-1(tm6311) V; csb-1(emc215[csb-1::AID:mScarlet3]) X |
| **HAL804** | uvs-1(emc97[uvs-1(tm6134)::AID::GFP]) V |
| **HAL812** | uvs-1(emc80) V; emcEx503[uvs-1::AID::GFP myo-2/myo-3::mCherry] |
| **HAL814** | uvs-1(tm6311) V; emcIs505[csb-1::FLAG::GFP elt-2::mCherry] |
| **HAL823** | uvs-1(emc510[S231A] (emc95[uvs-1::AID::GFP]) V |
| **HAL824** | uvs-1(tm6134) math-33(ok2974) V |
| **HAL825** | uvs-1(emc511[F339IDV342] (emc95[uvs-1::AID::GFP]) V |
| **HAL842** | uvs-1(emc80) V; emcEx522[uvs-1(tm6311)::AG XS myo-2/myo-3::mCherry] |
| **HAL843** | uvs-1(emc80) V; emcEx523[uvs-1(tm6134)::AG XS myo-2/myo-3::mCherry] |
| **HML1012** | cshIs140 [P(rps-28)::TIR1[F79G]::T2A::mCherry::his-11] II; ieSi58 [P(eft-3)::AID::GFP] IV |
| **OH16461** | ama-1(ot1037[GFP::3xFLAG::ama-1]) IV |
| **RB1426** | brd-1(ok1623) III |
| **RB2194** | math-33(ok2974) V |

### Supplementary Table 2. RT- and qPCR primers

| Primer | Sequence |
|---|---|
| ***uvs-1* forward** | ATGGATTCAATTGACAACTCC |
| ***uvs-1* reverse** | CCAGAATCTATGGCAATCTGC |
| **GFP forward** | GAAGGTGATGCAACATACGG |
| **GFP reverse** | CTTGACTTCAGCACGTGTCTTG |
| ***cdc-42* forward** | TCCACAGACCGACGTGTTTC |
| ***cdc-42* reverse** | AGGCACCCATTTTTCTCGGA |
| ***pmp-3* forward** | GTTCCCGTGTTCATCACTCAT |
| ***pmp-3* reverse** | ACACCGTCGAGAAGCTGTAGA |
